# Elucidation of the molecular mechanism of the breakage-fusion-bridge (BFB) cycle using a CRISPR-dCas9 cellular model

**DOI:** 10.1101/2024.04.03.587951

**Authors:** Manrose Singh, Kaitlin Raseley, Alexis M. Perez, Danny MacKenzie, Settapong T Kosiyatrakul, Sanket Desai, Noelle Batista, Navjot Guru, Katherine K. Loomba, Heba Z. Abid, Yilin Wang, Lars Udo-Bellner, Randy F. Stout, Carl L. Schildkraut, Ming Xiao, Dong Zhang

**Author notes:** Corresponding Authors: Dong Zhang, PhD, Department of Biomedical Sciences College of Osteopathic Medicine, New York Institute of Technology Rockefeller Building, Room 307, Northern Blvd, PO Box 8000, Old Westbury, NY 11568-8000. equal contribution (Manrose Singh, Kaitlin Raseley).

## Abstract

Chromosome instability (CIN) is frequently observed in many tumors. The breakage-fusion-bridge (BFB) cycle has been proposed to be one of the main drivers of CIN during tumorigenesis and tumor evolution. However, the detailed mechanisms for the individual steps of the BFB cycle warrants further investigation. Here, we demonstrated that a nuclease-dead Cas9 (dCas9) coupled with a telomere-specific single-guide RNA (sgTelo) can be used to model the BFB cycle. First, we showed that targeting dCas9 to telomeres using sgTelo impeded DNA replication at telomeres and induced a pronounced increase of replication stress and DNA damage. Using Single-Molecule Telomere Assay via Optical Mapping (SMTA-OM), we investigated the genome-wide features of telomeres in the dCas9/sgTelo cells and observed a dramatic increase of chromosome end fusions, including fusion/ITS+ and fusion/ITS-.Consistently, we also observed an increase in the formation of dicentric chromosomes, anaphase bridges, and intercellular telomeric chromosome bridges (ITCBs). Utilizing the dCas9/sgTelo system, we uncovered many novel molecular and structural features of the ITCB and demonstrated that multiple DNA repair pathways are implicated in the formation of ITCBs. Our studies shed new light on the molecular mechanisms of the BFB cycle, which will advance our understanding of tumorigenesis, tumor evolution, and drug resistance.

## Introduction

Genomic instability is one of the key hallmarks of cancer ^1-2^. Two main types of genomic instability are found in cancers: at the nucleotide level and at the chromosome level ^3^. The latter is referred to as chromosome instability (CIN), which has been observed in a wide variety of tumors ^4-5^. Though the underlying mechanisms of CIN are complex, the breakage-fusion-bridge (BFB) cycle is considered one of the major drivers of CIN ^4-6^. Dr. Barbara McClintock initially proposed the BFB cycle in the late 1930s by observing the cytogenetic changes in maize chromosomes after treatment with ionizing radiation (IR) ^7-8^. The current model for the BFB cycle proposes that the cycle starts with the formation of a dicentric chromosome, which is induced by one or more double-stranded DNA breaks (DSBs), frequently taking place at telomeres (Figure S1, Steps I and II). The piece(s) of chromosome detached from the main one can be temporarily compartmentalized in a micronuclei. If the dicentric chromosome is left unresolved and remains intact when the cell progress to anaphase, the dicentric chromosome then becomes an anaphase bridge when the two sets of chromosomes are pulled towards the opposing centrosomes by the mitotic spindles (Figure S1, Step III) ^9-11^. If the anaphase bridge fails to be resolved and persists during cytokinesis, it manifests as an intercellular chromosome bridge (Figure S1, Step IV) ^12-13^. The intercellular chromosome bridge needs to be severed before the generation of two fully separated daughter cells (Figure S1, Step V). If the broken chromosomes remain unrepaired in the daughter cells, they can potentially fuse with another broken chromosome thereby forming a new dicentric chromosome (Figure S1, Step VIa), and thus trigger the next BFB cycle. Alternatively, the broken chromosome can be healed via chromothripsis or some other mechanism(s) by re-incorporating the chromosome fragment protected in the micronuclei into the main chromosome and terminating the BFB cycle (Figure S1, Steps VIb and VII) ^5^. The BFB cycle plays a crucial role in promoting tumorigenesis and tumor evolution. For example, various solid tumors manifest many features of the BFB cycle occurrence ^6^. Most recently, using cultured cancer cell lines, Shoshani and colleagues showed that the BFB cycle and chromothripsis contribute to the rapid tumor evolution, leading to resistance to chemotherapy ^14^.

Though the concept of the BFB cycle was proposed more than eighty years ago ^7-8^, elucidation of the detailed mechanism of the BFB cycle has been challenging because naturally occurring dicentric chromosomes and resultant intercellular chromosome bridges in normal mammalian cells are rare events, which are too low for consistent detection, quantification, and characterization ^15-16^. In addition, the genetic loci where DSBs can occur and subsequently lead to the formation of a dicentric chromosome are difficult to identify. Intriguingly, in cells with dysfunctional telomeres, telomere fusion-induced dicentric chromosomes are drastically increased ^17-19^. Therefore, these cells have been used to investigate the nature of dicentric chromosome formation and the related DNA repair processes. For example, studies using yeast and mammalian cells with dysfunctional telomeres have implicated both canonical non-homologous end joining (cNHEJ) as well as alternative non-homologous end joining (altNHEJ) in promoting the formation of telomere fusion-induced dicentric chromosomes ^15, 18, 20-21^. AltNHEJ has been renamed microhomology-mediated end joining (MMEJ) or Polθ-mediated end joining (TMEJ) ^22^.

In particular, inducing telomere dysfunction by perturbing the function of the Shelterin complex has been an important tool of obtaining information on chromosome fusion and breakage. TRF2 is a key component of Shelterin that binds and protects the integrity of telomeres ^19^. Using an inducible dominant-negative TRF2 (TRF2-DN) model ^17^, two groups demonstrated recently that cytoplasmic nucleases such as TREX1 and the actomyosin-dependent physical stretching can facilitate the breakage of the intercellular chromosome bridges ^12-13^. Using the TRF2-DN cellular model, multiple groups have also implicated chromothripsis and kataegis as ways to heal the broken chromosome(s) in the daughter cells ^12-13, 23^. Additionally, Umbreit and colleagues have shown that depletion of a component of condesin using siRNA, treatment of cells with low-dose topoisomerase II inhibitor ICF-193, and CRISPR/Cas9-mediated cleavage of telomeres of chromosome 4 can also recapitulates certain features of the BFB cycle^13^.

Even though tremendous progress has been made in recent years to elucidate the detailed mechanism of the BFB cycle, many critical questions still need to be answered. For example, what type of genomic stress is capable of inducing the DSBs in Step I of the BFB cycle? To what extent are additional DSB repair pathways involved in the formation of dicentric chromosomes and intercellular chromosome bridges? Additionally, what is the molecular and structural nature of the intercellular chromosome bridge? Here, we first showed that targeting the nuclease-dead *Streptococcus pyogenes* Cas9 (dCas9) to telomeres with a telomeric single-guide (sg) RNA, or sgTelo, induces robust replication stress and DNA damage, specifically at telomeres. Remarkably, in the dCas9/sgTelo cells, we observed a drastic increase in the formation of chromosome end fusions, dicentric chromosomes, anaphase bridges, and intercellular telomeric chromosome bridges (ITCBs). Through this, we firmly established that the dCas9/sgTelo system can be used as a novel BFB cycle cellular model. Utilizing this model, we then investigated the mechanism of multiple steps of the BFB cycle. We not only confirmed the previous findings that cNHEJ promotes the formation of dicentric chromosomes, but further demonstrated that homology-dependent repair (HDR), including the break-induce replication (BIR), can also promote the formation of ITCBs. Most intriguingly, we found that TMEJ suppresses the formation of ITCB. Furthermore, we showed that when TMEJ is inhibited, cells upregulate HDR and downregulate cNHEJ to promote the formation of ITCB. Finally, we uncovered several new molecular and structural features of the ITCB through imaging. Collectively, our data shed new light on the detailed mechanisms of the BFB cycle, which will have an important impact on our understanding of tumorigenesis, tumor evolution, and drug resistance.

## Materials and Methods

### Cell lines and tissue culture

U2OS-derived dCas9/sgTelo and dCas9/sgNS cell lines were generously provided by Dr. Erik Sontheimer (University of Massachusetts Chan Medical School)^24^. The sequences for sgNS and sgTelo are: gAATCTCGCTTATATAACGAG and gTTAGGGTTAGGGTTAGGGTT respectively. All cells are grown in DMEM (Corning) supplemented with 10% fetal bovine serum (Bio-techne), penicillin (Corning), and streptomycin (Corning), at 37 °C in a humidified incubator with 5% CO_2._ To induce the expression of dCas9-mCherry-APEX2, cells are grown in media containing 2 μg/ml doxycycline (Thermo Fisher) and 250 nM Shield1 (Takara Bio) for 21 hours.

### Immunofluorescence

dCas9/sgTelo and dCas9/sgNS cells were seeded onto coverslips and induced with doxycycline and shield1 for 21 hours before fixation. Cells were either then fixed with 3% paraformaldehyde containing 2% sucrose for 10 min, followed by treatment with Triton X-100 solution on ice for 5 min, or treated with Triton X-100 solution on ice for 5 min then fixed with 3% paraformaldehyde containing 2% sucrose for 10 min (pre-extraction). Cells are stained with primary antibodies and the respective Alexa-488 (Invitrogen) and Alexa-546 (Invitrogen) conjugated secondary antibodies. All the antibodies used for IF are listed in **Supplemental Table 4**. SlowFade Gold DAPI (S36938) was used to stain the DNA and the DNA knots on ITCBs. Images were collected by using an Olympus upright Fluorescent Microscope or Axio Observer.Z1/7 (Zeiss Microimaging GmbH, Jena Germany). Images were processed using Adobe Photoshop or ZEN 2.5 pro.

### ImmunoFISH

Immunofluorescent and telomere fluorescent in situ hybridization was performed as previously described (Pan X., 2017). The telomere PNA probe (TelC) was purchased from PNA Bio (F1004). Images were collected by using an Axio Observer.Z1/7 (Zeiss Microimaging GmbH, Jena Germany).

### Single Molecule Analysis of Replicated DNA (SMARD)

The SMARD assay was performed essentially as described previously^25^. Briefly, dCas9/sgNS and dCas9/sgTelo cells were sequentially labeled with 30 μM IdU (4 hours) and 30 μM CldU (4 hours). DNA isolation and processing for SMARD were as described previously^25^. Following Pme I digestion, telomeric DNA fragments ranging from 160 to 200 kb were resolved by PFGE, identified by Southern blot, and then isolated. The Pme I-digested DNA was stretched on microscope slides coated with 3-aminopropyltriethoxysilane (Sigma). After stretching, the DNA was denatured in alkali-denaturing buffer (0.1 N NaOH in 70% ethanol and 0.1% β-mercaptoethanol) for 12 min and fixed by addition of 0.5% glutaraldehyde for 5 min. Telomeric DNA was identified by hybridization with a Biotin-OO-(CCCTAA)4 PNA probe (TelC, Biosynthesis) followed by incubation with Alexa Fluor 350-conjugated NeutrAvidin (Molecular Probes), followed by two rounds of incubation first with a biotinylated anti-avidin antibody (Vector) and then with the Alexa Fluor 350-conjugated NeutrAvidin. Incorporated halogenated nucleotides were detected with a mouse anti-IdU monoclonal antibody (BD) and a rat anti-CldU monoclonal antibody (Accurate) followed by Alexa Fluor 568-conjugated goat anti-mouse (Invitrogen) and Alexa Fluor 488-conjugated goat anti-rat secondary antibodies (Invitrogen). An antibody recognizing the single-stranded DNA was purchased from Millipore (anti-DNA antibody, clone 16–19, MAB3034).

### EdU Labeling for DNA Synthesis Detection

To visualize DNA synthesis, induced dCas9/sgTelo, and dCas9/sgNS cells on coverslips were incubated with 10mM EdU for 15min or 30 min prior to fixation. Cells on coverslip were cooled on ice rinsed once with 1x PBS, treated with pre-extraction buffer ( 0.1% Triton X-100, 20 mM HEPES-KOH pH 7.9, 50 mM NaCl, 3 mM MgCl2, 300 mM sucrose) for 5 min on ice, rinsed once with 1× PBS, fixed with 3% PFA and 2% sucrose for 15 min at room temperature and cold methanol for 10 min in −20°C. Cells were then permeabilized in PBS with 0.5%Triton X-100 for 3 min, and blocked with block solution (1×PBS containing 0.05% Tween-20 and 3% BSA) for 1h at RT. The Click-iT® EdU Imaging Kit (Cat. Nos C10337) was used to detect the EdU. Images were collected by using an Axio Observer.Z1/7 (Zeiss Microimaging GmbH, Jena Germany).

### Metaphase spread assay

For metaphase spread preparations, cells were plated at 1M cells per 10cm plate on Day 1. On Day 2, cells were induced with doxycycline and shield1 as previously described. For inhibitor studies, various DNA repair enzyme inhibitors were added at specified concentrations overnight (approximately 6 hours after induction). On Day 3, KaryoMAX Colcemid solution (Gibco, Catalogue No. 15212012) was added to the media to achieve a final concentration of 0.1 µg/mL for 2.5 hours. Cells were subsequently gently washed with PBS once, detached with 1mL of 0.25% trypsin and collected in media. Cells were centrifuged and pellet was washed with PBS and resuspended in 0.5mL PBS. PBS was removed and cells were resuspended in 5mL pre-warmed 37°C 0.075M KCl solution. Cells were incubated for 15 minutes at room temperature. 1mL of ice-cold fixative (3:1 volume of methanol: glacial acetic acid) was added dropwise and cells were incubated for an additional 10 minutes at room temperature. Cells were centrifuged for 5 minutes at 1000rpm and KCl solution was removed except for 0.5mL. Cells were resuspended in remaining KCl and additional 1mL of fixative was added while cells were vortexed at medium speed. An additional 8mL of fixative was added at high speed to ensure fixation. Cells were centrifuged again and resuspended in 200-300 µL of fresh fixative. Resuspended cells were dropped from several inches above the end of a dried tilted slide. Slides were airdried and mounted with DAPI. Metaphases were identified and imaged using a Axio Observer.Z1/7 (Zeiss Microimaging GmbH, Jena Germany) under 63x with oil immersion.

### Anaphase bridge assay

For analysis of anaphase bridges, cells were plated on coverslips one Day 1. On Day 2, cells were induced with doxycycline and shield1. On Day 3, Ro-3306 (Selleck, Catalog No. S7747) was added to the media for a final concentration of 7µM and incubated at 37°C for 6 hours. After incubation, cells were washed three times with pre-warmed media and the specified DNA repair enzyme inhibitors were added for 1 hour and 15 minutes. Coverslips were then washed with PBS once and fixed at room temperature in 3% paraformaldehyde and 2% sucrose solution. Cells were washed again and mounted onto slides using DAPI solution. Anaphases were identified and imaged using a Axio Observer.Z1/7 (Zeiss Microimaging GmbH, Jena Germany) under 63x with oil immersion.

### Time-lapse imaging of live cells

Time-lapse imaging of live cells was done for 25 hours with a time interval of 30 min between time points on a fully automated DVCore microscope (Leica-microsystems), using a 60x oil objective (N.A. 1.42) a chamber heated to 37°C and a Cascade2 EMCCD camera.”

### siRNA Transfection

Cells are transfected twice on consecutive days using 50 nM siRNA with RNAiMAX (Invitrogen) following the protocol recommended by the manufacturer. The sequence and order information for all the siRNA are listed in **Supplemental Table 5**.

### Long-term viability assay

4000 cells/per well are seeded in 12 well plates. Cells are induced the following day with media containing doxycycline and shield1. After 7-10 days of growth, cells are fixed using a methanol/acetic acid solution and stained with 1% crystal violet solution.

### Single-molecule telomere assay via optical mapping (SMTA-OM)

#### DNA extraction and purification

Cells were first embedded in gel plugs with approximately 1 million cells per gel plug (BioRad no. 170-3592). The high molecular weight genomic DNA was extracted and purified using the Bionano Genomics SP kit.

#### DNA Labeling

Following the manufacturer’s instructions, a DLS labeling kit (Bionano Genomics) labeled 750 ng of genomic DNA. A labeling mix comprised Direct Labeling Enzyme 1 (DLE-1), 1X DLS reaction buffer, and DL green fluorophore-labeled nucleotide mix. The labeling mix was added to the genomic DNA, gently mixed via a wide bore pipette, and then incubated at 37°C for 2 hours. Following the incubation, membrane dialysis was used to remove the unwanted fluorescent dyes, proteins, and salts. Membrane dialysis was done at room temperature for approximately 2 hours in the dark to protect it from the light. Dialyzed DNA was then recovered using a 100 nm hydrophilic membrane (EMD Millipore, VCWP04700) and quantified with a Qubit device.

The second part of the labeling procedure focuses on labeling the telomeres. Guide RNA (gRNA) consisting of 0.5 μM tracrRNA (IDT) and 50 μM crRNA was gently mixed via pipetting up and down and annealed on ice for 30 minutes. 25 pmol gRNA was incubated with 200 ng Cas9D10A nicking enzyme and 1X NE Buffer 3.1 at 37°C for 15 minutes. Next, 300 ng of the previously DLE-1 labeled genomic DNA was added to the Cas9D10A mixture for the one-hour nicking reaction at 37°C. After nicking, 200 nM red fluorophore-labeled nucleotides (ATTO647-dUTP, dATP, dGTP, dCTP) were incorporated into telomeres by 5U Taq DNA polymerase at 72°C for one hour in 1X Thermopol buffer (New England Biolabs). The nick-labeled sample was treated with proteinase K (QIAGEN) at 50°C for 30 min. Finally, to prepare the DNA for nanochannels, a staining mix of flow buffer, DTT, and YOYO-1 (a blue-colored dye that labels the backbone of DNA, Bionano Genomics DLS kit) was prepared according to the manufacturer’s instruction before adding it to the sample. The sample combined with the staining mix was incubated at room temperature overnight.

#### Imaging with Saphyr

A Bionano Saphyr G1.2 chip was loaded with the labeled DNA sample and imaged using a “dual-labeled sample” scheme incorporated in the Saphyr software. Images were taken in the following order: first, the red fluorescent labeling with the 637 nm laser, then the green fluorescent labeling with the 532 nm laser, and finally, the blue fluorescent DNA backbone YOYO-1 staining with the 473 nm laser.

#### Genome Assembly

*De novo* genome assembly was done using the software developed by Bionano Genomics. Consensus maps were generated by de novo assembling individual DNA molecules followed by alignment to the hg38 human reference genome.

#### Telomere analysis

CMAP, XMAP, and BNX files were generated after *de novo* genome assembly. The CMAP files contain both the red and the green label information. The XMAP files contain only the information on the green labels. Molecules matching the expected green labeling patterns were extracted from the BNX and CMAP files. Raw molecule images were extracted using in-house software based on their locations in the nanochannels. The red telomere labels were easily distinguishable from the green labels in the subtelomeric region. Telomeres were then analyzed and measured using the ImageJ software. The ferret diameter tool was used to capture the length of telomeres in pixels and intensities before converting all the measurements to kilobase (kb) following the previously established procedure^26-27^.

### Inhibitor treatment assay

All small molecule inhibitors used in this study are listed in the Supplemental Table 6. For all small molecule inhibitor treatments, cells are plated on day 1. On day 2, the dcas9/sgTelo expression is first induced with doxycycline/shield1 for 6 hours. These cells are then treated with the chosen concentrations of each inhibitor for 15 hours. After 21 hours from the initial induction, cells were fixed, stained, and imaged.

### Immunoblotting

Cells are collected and lysed directly in a sample buffer, followed by sonication for 30 seconds. Equal volumes of lysate were loaded for immunoblotting. Antibodies used for immunoblotting: Actin (Santa Cruz, sc-1616); BLM (Bethyl, A300-110A, and A300-120A), BRCA1 (EMD/Calbiochem, OP92); BRCA2 (EMD/Calbiochem, OP95), Rad51 (Santa Cruz, sc-8349); Rad52 (Santa Cruz, sc-365341); POLD3 (Abcam, 182564).

### Immunostaining and Super-Resolution Imaging (STORM)

dCas9/sgTelo cells were seeded at a density of 3.5 x 10^5^ on coverslips in 6-well plates. The next day, cells were induced for 21 hours with shield1 and doxycycline. Immediately after, cells were fixed with 3% PFA and 2% sucrose solution, washed with PBS, and permeabilized in a Triton X-100 solution. Primary antibody staining was done with RPA32-pS4pS8 (Bethyl, A300-245A), TRF1 (abcam, ab10579) and BLM (Bethyl, A300-110A). Secondary staining was done with anti-rabbit Alex Fluor 647 (Thermo Fisher, A32733), and the endogenous mCherry signal was boosted with anti-alpaca Alexa Fluor 568 RFP-booster (Chromotrek, rb2AF586). Unbound antibodies were washed off, cells were treated with 2% PFA and 3% sucrose solution for 10 minutes and again washed with PBS.

Super-resolution image acquisition was done with a Nanoimager super-resolution microscope (Oxford Nanoimaging Ltd., Oxford, UK), which was used with an Olympus 1.4NA 100× oil immersion super apochromatic objective. Initial calibration was done using 0.1 μM Tetraspeck beads (Thermo Fisher). Cells were flushed with blinking induction buffer containing glucose, glucose oxidase, catalase, and β-mercaptoethanol prior to every imaging session. Excitation of Alexa Fluor 647 and RFP-boosted mCherry fluorophores was done using lasers set at 640 nm and 568 nm, respectively. Initially, the 568 nm laser was used at 3-4% power for an overview scan of a partial region of the coverslip to identify bridges. Following identifying a bridge of interest, multi-image acquisition was set up to take images at 30 ms/frame for each wavelength as a series of 5000 or 10000 frames. Excitation at 640 nm was done first, followed by at 568 nm to limit potential photobleaching. Following image acquisition, initial 50-100 frames were filtered out, and the remaining were used for analysis.

## Results

### Targeting dCas9 to telomeres induces replication stress, DNA damage, and micronuclei formation

Previously, dCas9 was shown to be a targetable roadblock for DNA replication in vitro ^28^. In a separate study, Sontheimer and his colleagues developed a doxycycline (Dox)-inducible system in U2OS cells to identify proteins near a well-defined genomic locus called C-BERST (d<u>C</u>as9-APEX2 <u>b</u>iotinylation at genomic <u>e</u>lements by <u>r</u>estricted <u>s</u>patial tagging, Fig 1A) ^24^. They designed two sgRNAs, sgAlpha, which targets the dCas9-APEX2 to centromeres, and sgTelo, which targets the dCas9-APEX2 to telomeres. A sgRNA targeting non-specific DNA sequences, or sgNS, was used as the negative control. They identified many well-known centromere and telomere-associated proteins through mass spectrometry. Intriguingly, they also identified many DNA damage response (DDR) proteins at both centromeres and telomeres. Among them are BLM, BRCA1, and FANCM, which we have shown previously to suppress the replication stress at Alternative Lengthening of Telomere (ALT) telomeres ^29^. Intrigued by the results of this study, we hypothesized that targeting the dCas9-APEX2 protein to telomeres impedes the progression of the replisome, leading to stalling of the replication forks at telomeres and activation of the replication stress response and DNA damage response (DDR) pathways. An additional and useful feature of the C-BERST system is the inclusion of a mCherry marker within the dCas9-APEX2 fusion (Fig 1A), which can be conveniently used to visualize the locations of dCas9/sgRNA in both fixed and live cells.

**Figure 1.**
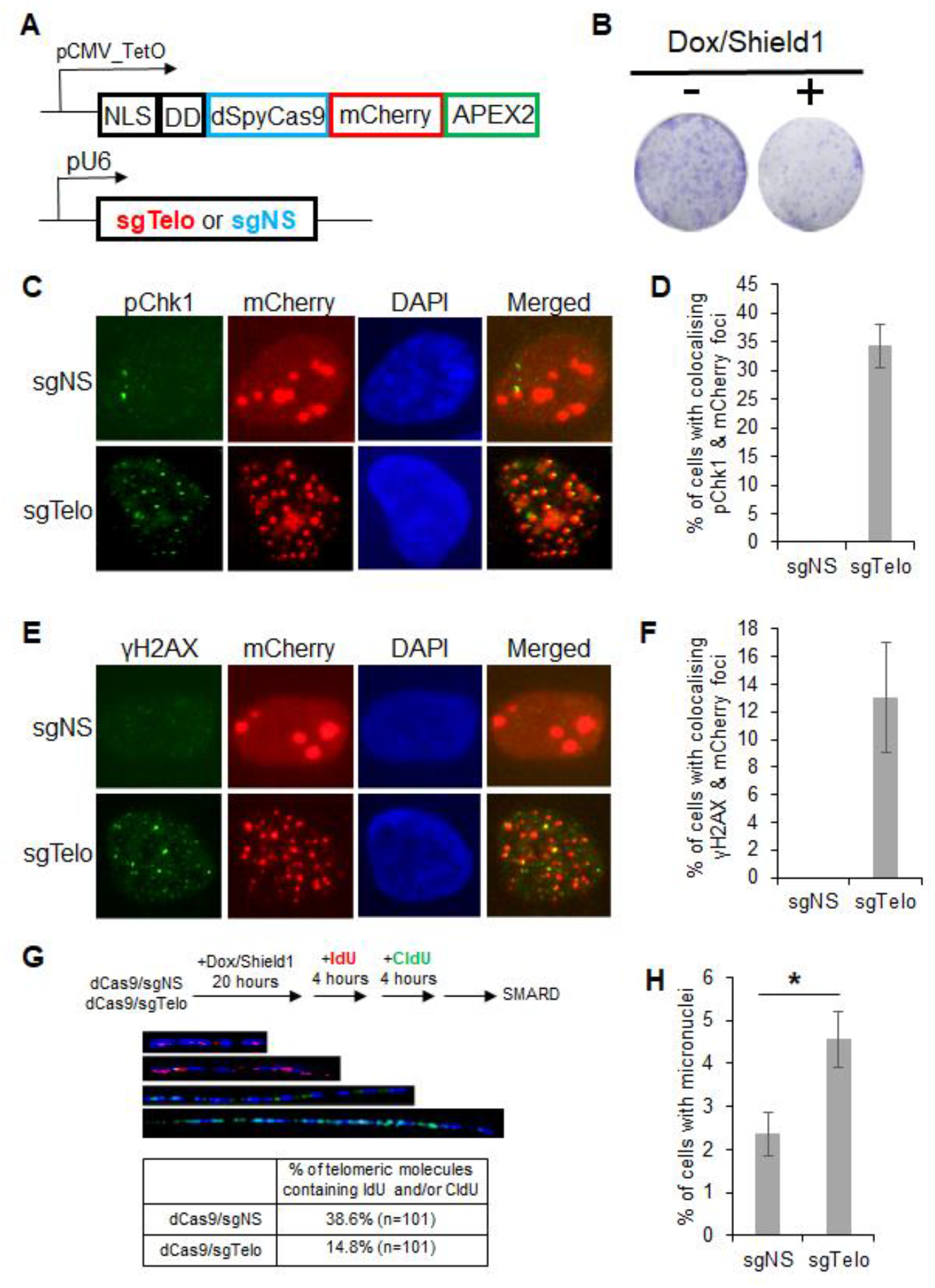
Targeting dCas9 to telomeres blocks DNA replication and induces pronounced replication stress and DNA damage. (**A**) A diagram of the inducible targeting of dCas9-mCherry-APEX2 specifically to telomeres. NLS: nuclear localization signal; DD: Shield1-repressible degradation domain. The expression of dCas9-mCherry-APEX2 is controlled by a doxycycline (Dox) inducible promoter (pCMV-TetO). dCas9-mCherry-APEX2 is further stabilized by adding Shield1, which inactivates the DD domain. **sgNS**: a non-specific single-guide RNA; **sgTelo**: a telomeric single-guide RNA. **(B)** Long-term viability assay comparing non-induced (-) and induced (+, with the addition of both Dox and Shield1) dCas9/sgTelo cells. **(C to F)** 21 hours after the addition of Dox and Shield1, dCas9/sgNS and dCas9/sgTelo cells were fixed and stained with antibodies recognizing either phosphorylated Chk1 (labeled as pChk1 in **C** and **D**) or γH2AX (**E** and **F**). All nuclei were also stained with DAPI. More than 200 cells were counted. Error bars are standard deviations from three independent experiments. (**G**) dCas9/sgTelo reduces the replication efficiency of telomeric DNA. dCas9/sgNS and dCas9/sgTelo cells were first treated with Dox and Shield1 for 20 hours and then sequentially pulse-labeled with IdU and CldU for 4 hours each. SMARD was performed on telomeric DNA PmeI fragments isolated from these cells as detailed in STAR METHODS, with fragments ranging from 160 to 200 kb. The telomeric DNA molecules of variable lengths were identified by FISH using telomeric PNA probes (TelC, blue). The incorporated IdU and CldU were detected by indirect immunofluorescence with antibodies recognizing IdU (red) and CldU (green). Four representative telomeric molecules are shown. (**H**) Increased micronuclei formation in dCas9/sgTelo cells. More than 1000 cells were counted. Error bars are standard deviations from two independent experiments. Standard two-tailed Student’s t-test: ^*^: p < 0.05.

To test our hypothesis, we first confirmed that a 21-hour treatment of dCas9/sgTelo cells with Dox and Shield1 led to an increase of mCherry foci that colocalize with both TelC foci and TRF1 foci (Figure S2A). TelC is a telomere-specific PNA probe that detects telomeric DNA through *in situ* hybridization ^29^. TRF1 is part of the Shelterin complex and is also widely used as a marker for telomeres ^19^. Therefore, mCherry can also be used as a reliable telomere marker in the dCas9/sgTelo cells. Targeting dCas9-mCherry-APEX2, abbreviated as dCas9, to telomeres compromised the long-term viability of the dCas9/sgTelo cells (Fig 1B) and induced robust replication stress and DNA damage at telomeres as indicated by the increased colocalization of mCherry foci with phosphorylated Serine-345 of Chk1 (pChk1) foci and with γH2AX foci, respectively (Figs 1C to 1 F). The patchy red signals seen in the dCas9/sgNS cells are likely due to the aggregation of mCherry in the nuclei (Figs 1C and 1E). In addition to pChk1 and γH2AX, many other well-known DDR proteins, including BLM and RPA32, can also be detected in the mCherry foci (Figs S2B to S2G). To examine whether dCas9/sgTelo directly affects DNA replication at telomeres, we performed the Single Molecule Analysis of Replication DNA (SMARD) assay ^29^. Indeed, we observed a 2.6-fold decrease in the replication efficiency of telomeric DNA in the dCas9/sgTelo cells compared to dCas9/sgNS cells (Fig 1G). Consistent with the heightened replication stress and increased DNA damage at the telomeres of the dCas9/sgTelo cells, we also observed a two-fold increase in the micronuclei formation (Fig 1H).

Taken together, our data strongly indicate that targeting dCas9 to telomeres physically impedes the progression of DNA replication machinery at telomeres. This leads to stalled replication forks, the activation of a robust replication stress response, increased DNA damage, and micronuclei formation and eventually causes decreased cell viability.

### dCas9/sgTelo promotes the formation of chromosome end fusions

Having established that targeting dCas9 to telomeres induces a robust replication stress response and DNA damage, we sought to thoroughly characterize the genome-wide changes at telomeres and chromosome ends in the dCas9/sgTelo cells using the recently developed <u>S</u>ingle-<u>M</u>olecule <u>T</u>elomere <u>A</u>ssay via <u>O</u>ptical <u>M</u>apping (SMTA-OM) ^26-27, 30-31^. The principle of three-color SMTA-OM and some of its readouts are illustrated in Figure S3. Briefly, long DNA fibers are carefully extracted and purified from cultured cells. The long DNA fibers (>150 kb) are first tagged with a green fluorescent label by the DLE-1 enzyme (Figure S3A) ^32^. Telomeres are then specifically labeled with a red fluorescent label by CRISPR/Cas9 (Figure S3A) ^26^. DNA backbones are stained with a blue DNA dye, YOYO-1. The DNA fibers are linearized through nanochannels and imaged. The following telomere/chromosome end features can be obtained from a single SMTA-OM assay (Figs 2A and 2B, Figure S3B). (1) End telomere (End Tel) refers to a DNA molecule that is embedded in a blue labeled DNA molecule and has a red signal (telomere) at the end of a specific chromosome arm, which is identified by the pattern of green labels matched to the hg38 reference genome. The intensity of the red signal is used to measure the length of a telomere of a specific chromosome arm. (2) Telomere-free end (TFE) refers to a specific chromosome arm without any detectable red signal at its end. (3) Fusion/ITS+, or ITS+: one side of a red signal is attached to an identifiable chromosome arm, while the other side of the red signal is also attached to a green-labeled DNA fragment. However, because the latter DNA fragment is too short, it cannot be unambiguously assigned to any specific chromosome arm based on the pattern of the green labels. (4) Fusion/ITS-, or ITS-: one portion of the green-labeled DNA fragment can be assigned to a specific chromosome arm based on the pattern of the green labels, while the other portion cannot be assigned to any chromosome arm again because the DNA fragment is too short. Meanwhile, there is no detectable telomeric signal (RED) in between the two DNA fragments. Compared to other widely used telomere assays ^33^, such as telomere restriction fragment assay (TRF), quantitative fluorescent *in situ* hybridization (Q-FISH), and STELA, the SMTA-OM offers several unique advantages. (1) SMTA-OM can measure a broad telomere length range, from 100 bp to more than 100 kb. (2) SMTA-OM can unambiguously assign a telomere to a specific chromosome arm. Therefore, it can monitor genome-wide telomere changes at the specific chromosome arm level and the single-molecule level. (3) SMTA-OM can also detect the TFE of a specific chromosome arm induced by DNA damage in the subtelomeres. (4) SMTA-OM can unambiguously assign one portion of the fused chromosome in ITS+ or ITS-to a specific chromosome arm.

**Figure 2.**
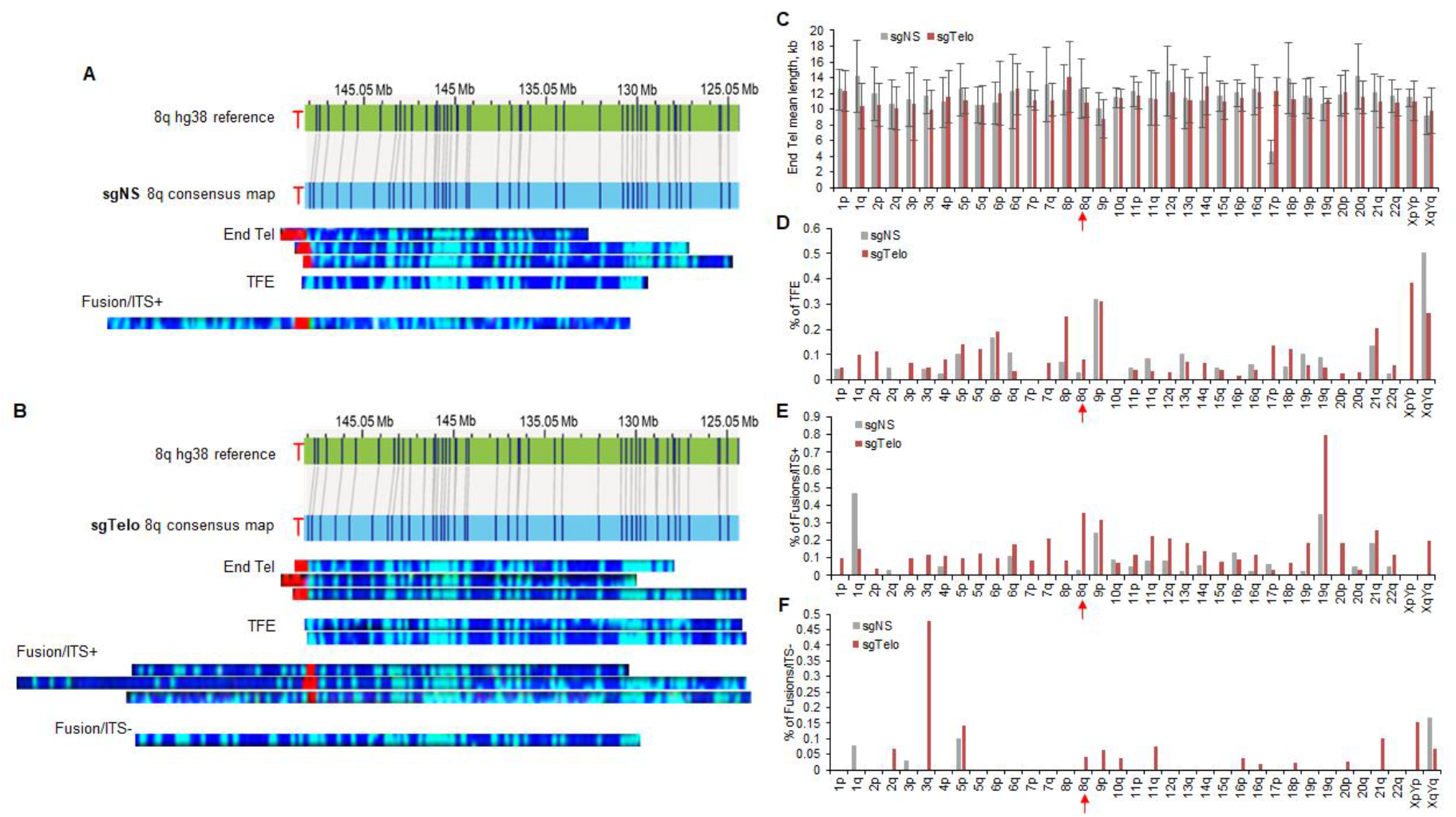
Genome-wide analysis of telomeres/chromosomes ends at the individual chromosome arm level. dCas9/sgNS and dCas9/sgTelo cells were treated with doxycycline and Shield1 for 21 hours and then were used for SMTA-OM analysis. (**A** and **B**) representative optical mapping images of chromosome arm 8q from the dCas9/sgNS cells (**A**) and dCas9/sgTelo cells (**B**). (**C** to **F**) The SMTA-OM profiles of 35 chromosome arms, including End Tel mean length (**C**), percentage of TFE (**D**), percentage of fusion/ITS+ (**E**), and percentage of fusion/ITS- (**F**). Red arrows highlight chromosome arm 8q.

Using the SMTA-OM, we comprehensively characterized the aforementioned telomere/chromosome end features of 35 chromosome arms by imaging approximately 30 molecules per arm for both dCas9/sgTelo and dCas9/sgNS cells. The optical mapping images of two representative chromosome arms, 8q and 3q, are shown in Figs 2A and 2B, and Figures S4 and S5. The raw measurements of the 35 chromosome arms are included in the Supplemental Table 1. The number of single molecules for each arm and calculated mean lengths, standard deviations, and frequencies of end telomere, TFE, ITS+, and ITS-events per arm from both samples are summarized in Supplementary Table 2.

**Table 1:**
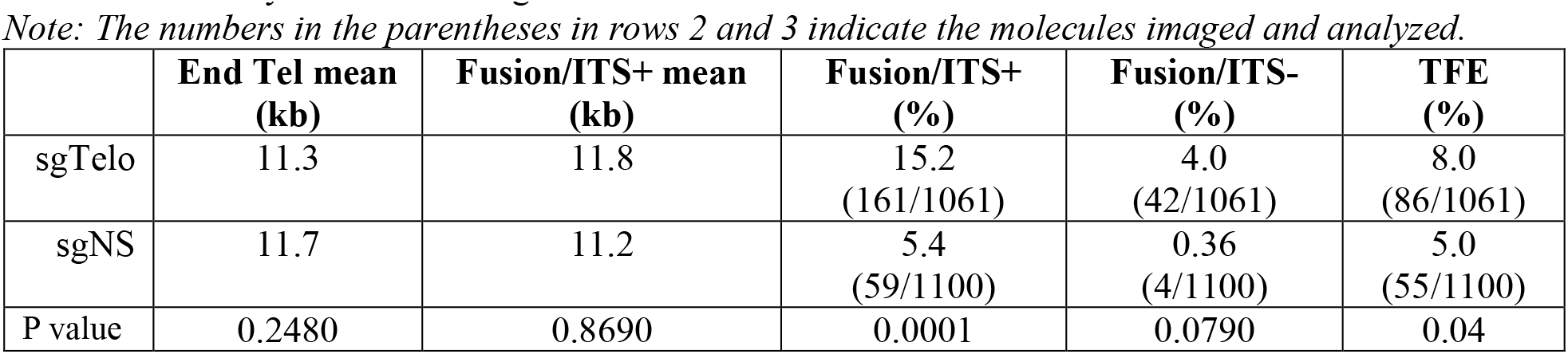
Summary of overall changes at telomeres/chromosome ends of different chromosome arms.

Regarding the length of End Tel, we observed significant changes for particular chromosome arms (Fig 2C, Figure S6, and Table 1). For example, the length of telomeres of 14q showed a 17% increase (13 kb for dCas9/sgTelo and 11.1 kb for dCas9/sgNS), while those of 20q showed an 19% decrease (11.5 kb for dCas9/sgTelo and 14.2 kb for dCas9/sgNS), suggesting there are chromosome arm-specific changes. However, when considering all the chromosome arms examined together, the overall telomere length change of End Tel is statically insignificant between dCas9/sgTelo and dCas9/sgNS (11.3 kb for dCas9/sgTelo and 11.7 kb for dCas9/sgNS, Figure S6 and Table 1). Regarding the TFEs, we observed a 60% overall increase in frequency within dCas9/sgTelo cells compared to the control (Fig 2D and Figure S7, Table 1), suggesting that many chromosome arms in dCas9/sgTelo cells have lost most of their telomeres.

When examining telomere fusions, we observed a 2.8-fold and a 11.1-fold increase for the fusion/ITS+ (Fig 2E and Figure S8, Table 1) and fusion/ITS-(Fig 2F and Figure S9, Table 1) in the dCas9/sgTelo cells compared to the dCas9/sgNS cells, respectively. Since fusion/ITS-is potentially formed from two TFEs, the drastic increase of fusion/ITS-suggests that certain chromosome arms in the dCas9/sgTelo cells have lost most of their telomeres, which is consistent with the increase of TFEs. Similar to the End Tel findings, there are notable changes for different chromosome arms. For example, the fusion/ITS+ increased by 4.9-fold for 8q (Figure S8). For 3q, the fusion/ITS-increased from 0% in the dCas9/sgNS cells to 47.6% in the dCas9/sgTelo cells (Figure S9). There are no overall changes in the mean length of telomeres in fusion/ITS+ (Figure S10 and Table 1).

In summary, the SMTA-OM analysis demonstrates that in the dCas9/sgTelo cells, the heightened replication stress at telomeres induces dramatic telomeric and subtelomeric DNA damage, leading to increased TFEs, fusion/ITS+, and fusion/ITS-. Most importantly, we also observed pronounced chromosome arm-specific changes for all the SMTA-OM readouts, indicating that telomeres of different chromosome arms respond to replication stress differently. These results underscore the importance of monitoring the telomere/chromosome end changes at the individual chromosome arm level.

### dCas9/sgTelo promotes the formation of dicentric chromosomes and anaphase bridges

The drastic increase of fusion/ITS+ and fusion/ITS-observed from the SMTA-OM analysis suggests that they are likely derived from different types of dicentric chromosomes induced by DSBs generated within telomeres, or at the subtelomeres, or the combination of both. We performed metaphase spread assays to monitor the changes in dicentric chromosomes frequency directly. As shown in Figs 3A and 3B, we observed a 3-fold increase of dicentric chromosomes in the dCas9/sgTelo cells compared to the dCas9/sgNS cells. Consistent with the increase of dicentric chromosomes, we also observed a 2-fold increase of anaphase bridges in the dCas9/sgTelo cells compared to the dCas9/sgNS cells (Figs 3C and 3D),suggesting that the Majority of the dicentric chromosomes remain unresolved and persist into the anaphase.

**Figure 3.**
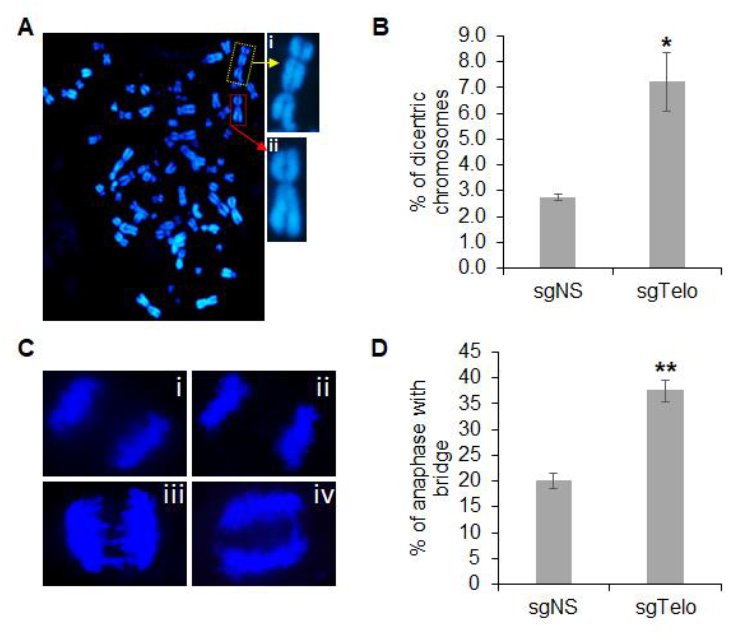
Targeting dCas9 to telomeres dramatically increases dicentric chromosomes and anaphase bridges. (**A** and **B**) dCas9/sgNS and dCas9/sgTelo cells were treated with doxycycline and Shield1 for 21 hours and then were used for metaphase spread analysis. **A**-i: The representative image of a dicentric chromosome. **A**-ii: the representative image of a normal monocentric chromosome. More than 2500 metaphase chromosomes were analyzed and qualified per condition. (**C** and **D**) dCas9/sgNS and dCas9/sgTelo cells were treated with doxycycline and Shield1 for 21 hours and then were used for anaphase bridge analysis. Minimum 50 anaphase cells were analyzed and quantified. Error bars are standard deviations from three independent experiments. Standard two-tailed Student’s t-test: ^*^: p < 0.05; ^**^: p < 0.01.

### dCas9/sgTelo promotes the formation of intercellular telomeric chromosome bridges

Next, we investigated the cellular consequences of the drastically increased chromosome fusions, dicentric chromosomes, and anaphase bridges in the dCas9/sgTelo cells. We reasoned that without their successful resolution, the fused chromosomes could eventually lead to the formation of intercellular chromosome bridges during cytokinesis (Step IV, Figure S1). Indeed, after imaging thousands of cells, we detected intercellular telomeric chromosome bridges (ITCBs), manifesting as two interphase cells with de-condensed chromosomes connected by a red/mCherry positive bridge, in ∼3% of the dCas9/sgTelo cells (Figs 4A and 4B). Previously, a non-specific double-stranded DNA binding protein, Barrier-to-Autointegration Factor (BAF), has been used as a marker for the intercellular chromosome bridges ^12-13^. Every ITCB we identified was also positive for BAF (Fig 4A-i). And when we stained the ITCBs with the TelC probe, all the mCherry bridges were also positive for TelC, confirming that the ITCBs indeed consisted of telomeric DNA. When the BAF marker and TelC probes were used to stain the control sgNS cells, we identified BAF+ bridges at a frequency of 3%, but none of these bridges were TelC-positive or mCherry-positive (Fig 4B). Collectively, our data indicates that the formation of ITCB is a rare phenomenon in the non-stressed control cells. However, the induction of the dCas9/sgTelo triggers the formation of predominantly telomere-mediated fusions and subsequently ITCBs with telomeric DNA.

**Figure 4.**
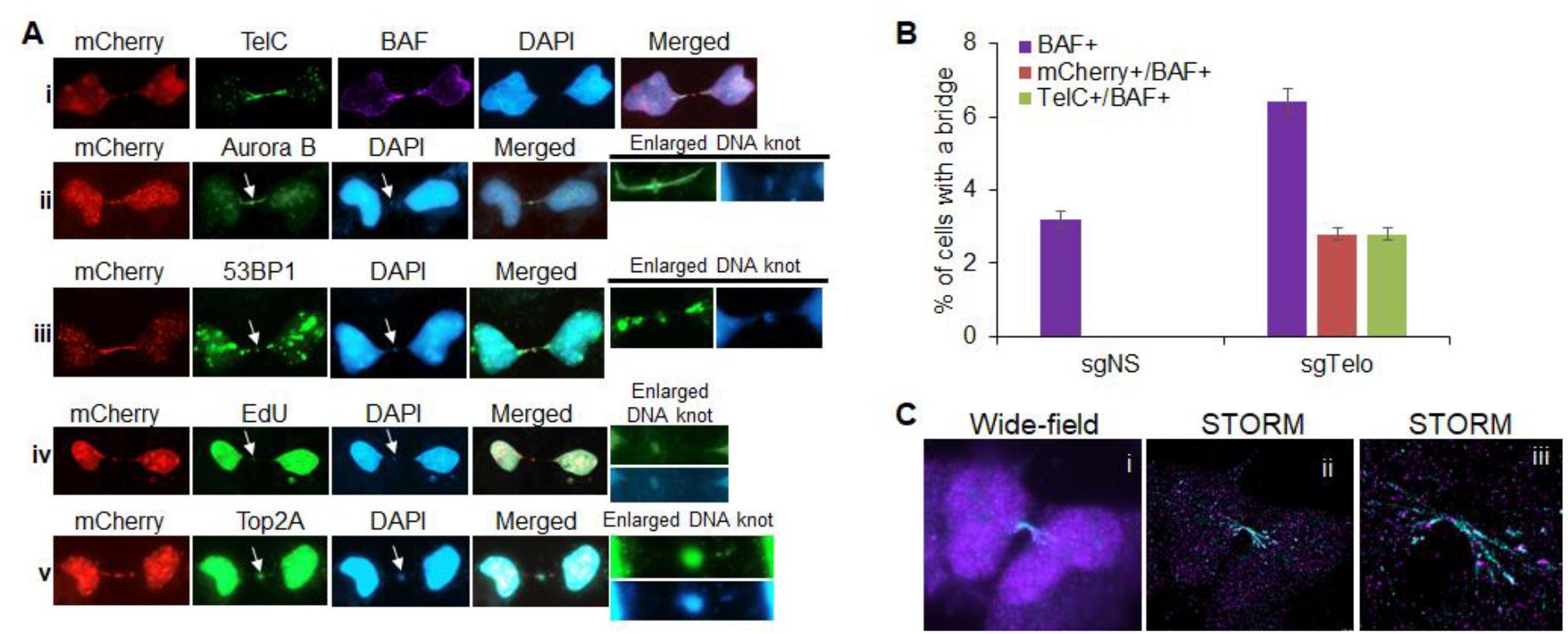
Targeting dCas9 to telomeres induces a drastic increase in intercellular telomeric chromosome bridges (ITCBs). (**A**) dCas9/sgTelo cells were treated with doxycycline and Shield1 for 21 hours. Cells were then fixed and stained with either a PNA probe recognizing telomeres (TelC) or antibodies recognizing different proteins as indicated. For the EdU pulse labeling image (**A**-iv), EdU was added to the growth medium 30 min before fixation. (**B**) More than 3000 cells were analyzed and quantified. (**C**) dCas9/sgTelo cells were treated with doxycycline and Shield1 for 21 hours. Cells were then fixed, stained with antibodies recognizing mCherry, and were imaged with the STORM. (**C**-i) wide-field imaging of an ITCB. (**C**-ii) STORM imaging of the same ITCB. (**C**-iii) Zooming image of the same ITCB.

We then sought to carefully characterize the structure of the ITCBs and delineate staining patterns at the bridge body and the bridge bases. We observed that intriguingly, on approximately 86% of the ITCBs, we can detect a more intense DAPI-positive punctate structure in the middle of the ITCB, which we denote as a “DNA knot” (Figs 4A-ii to 4A-v, Figure S11). We also noted that approximately 83% of the BAF-positive intercellular chromosome bridges in the sgNS cells also contain a DNA knot, suggesting that the formation of the DNA knot is independent of whether the bridge is induced by telomeric fusion or non-telomeric fusion. Given the location of the DNA knot occurring between two daughter cells, we hypothesized this could be the location of the midbody. Indeed, 100% of the DNA knots are stained positive for Aurora B (Fig 4A-ii), a commonly used marker for the midbody, indicating that the DNA knots are likely part of the midbodies. We also stained the ITCBs with an antibody recognizing 53BP1, a master regulator of DDR and DSB repair pathway choices ^34-35^. We not only detected 53BP1 at 70% of the DNA knots but also at the 100% of the bridge bases and 56% of the bridge bodies (Fig 4A-iii and Figure S11). To test whether there is an active DNA synthesis at the DNA knots, we pulse-labeled dCas9/sgTelo cells with EdU for either 15 min or 30 min prior to fixation (Fig 4-iv and Figure S12). We observed positive EdU signals at 31% and 38% of the DNA knots, respectively, indicating active DNA synthesis in one third of the DNA knots. In addition, more than half of the DNA knots are also stained positive for Top2A, suggesting that there may be over-twisted or catenated DNA in the DNA knots (Fig 4A-v and Figure S13).

In light of the discovery of DNA knots and the interesting staining patterns of 53BP1 and Top2A, we surveyed a panel of DDR proteins in the ITCB-forming cells. The DDR proteins surveyed so far manifest three distinct staining patterns: (1) DDR proteins, such as 53BP1, RPA32, cGAS, and FANCI, stain positive of the bridge body (Fig 4A-ii, Figures S14A to S14C and S14N). Positive staining of RPA32 on the bodies of ITCBs suggests that there are single-stranded DNA (ssDNA) regions on the ITCBs. (2) DDR proteins, such as 53BP1, γH2AX, and pChk1 stain positive of one or both bridge bases (Fig 4A-iii and Figures S14D and S14E, S14N). (3) DDR proteins, including 53BP1, FANCM, BLM, and Rad51, localize in the middle of the bridges (Fig 4A-iii, Figures S14F to S14M, and S14N).

To gain further structural insights into the ITCBs, we also performed <u>St</u>ochastic <u>O</u>ptical <u>R</u>econstruction <u>M</u>icroscopy (STORM). We observed that most ITCBs are branched and have a similar number of branches on both sides of the bridges (Fig 4C). We speculate that these branches reflect the number of dicentric chromosomes in the ITCBs. The average branches per ITCB are 2.846 (Figure S15A and Supplemental Table 3), with an average length of 6.353 μm (Figure S15B and Supplemental Table 3) and an average width of 0.285 μm (Figure S15C and Supplemental Table 3). These data suggest that ITCBs may contain bundled dicentric chromosomes.

### Formation of an ITCB in the dCas9/sgTelo cells delays the completion of cytokinesis

Next, we investigated the effects of the ITCBs on cell cycle progression in live cells by performing a time-lapse experiment. When the expression of mCherry is relatively low, the cell progresses through mitosis and cytokinesis without delay. It usually takes less than 30 min for all the chromosomes to align at the metaphase plate and complete cytokinesis and produce two daughter cells (Fig 5A, Figures S16A and S16B, and Supplemental Movie 1). In stark contrast, in the cell that eventually forms an ITCB, the time between metaphase and completion of cytokinesis increases to between 90 min and 120 min (Fig 5B, Figure S16C, and Supplemental Movie 2). This suggests that the presence of an ITCB triggers the activation of the abscission checkpoint, thereby delaying the completion of cytokinesis to give the cell more time to resolve the ITCB. We also noted that many mCherry-positive cells failed to progress through the different stages of mitosis and eventually committed to cell death, vis-à-vis mitotic catastrophe (Fig 5C, Figure S16D, and Supplemental Movie 1).

**Figure 5.**
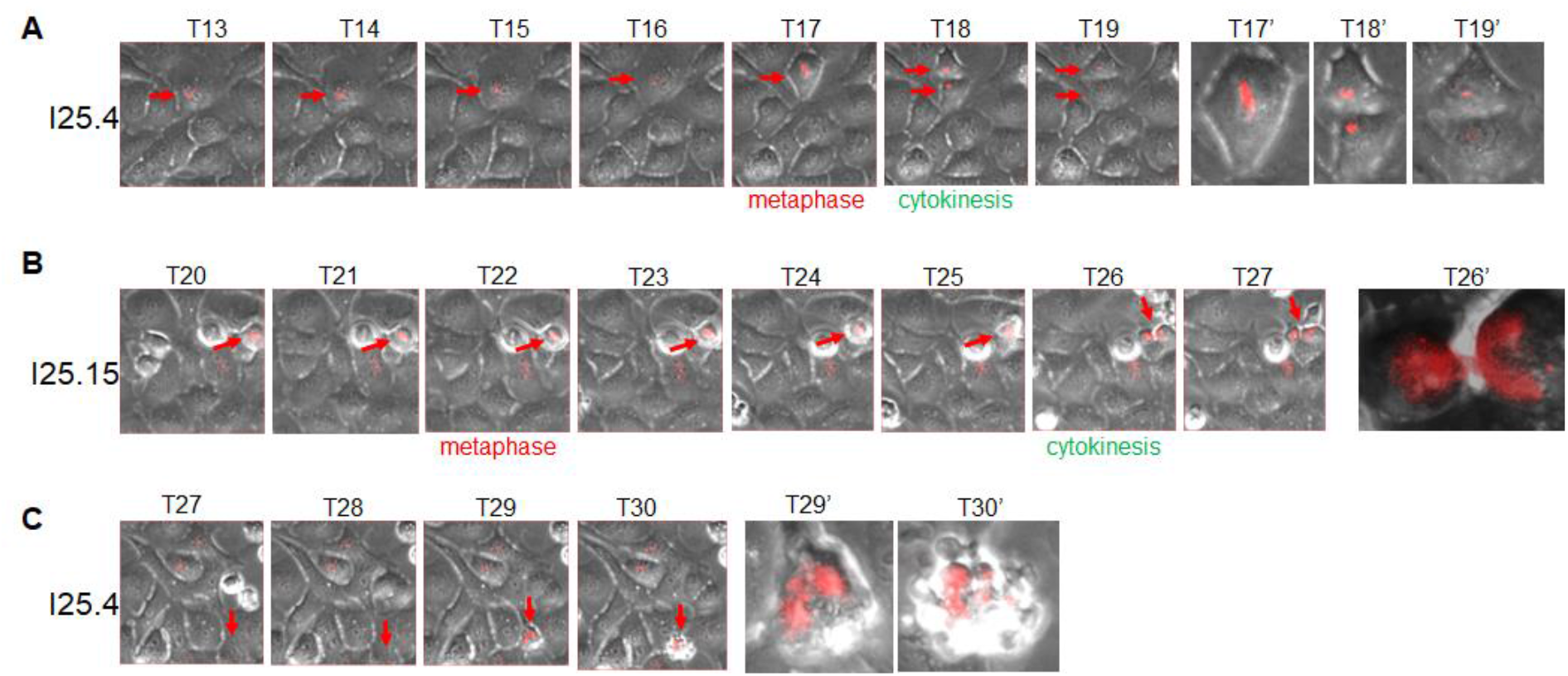
Time-lapse analysis of live dCas9/sgTelo cells. Asynchronous dCas9/sgTelo cells were pre-treated with doxycycline and Shield1 for 8 hours. The cells were kept in doxycycline and Shield1 and monitored using a fully automated DVCore microscope (Leica-microsystems). Images were acquired every 30 minutes for 25 hours as indicated by the Tn. Tn’ is the enlarged view of the corresponding Tn. (**A**) The red arrows indicate a cell that expressed a low level of mCherry and manifested relatively normal cell cycle progression. (**B**) The red arrows indicate a cell that formed the intercellular telomeric chromosome bridge at T26. (**C**) The red arrows indicate a cell that underwent a mitotic catastrophe.

In summary, we have shown that: (1) targeting dCas9/sgTelo to telomeres induces robust replication stress, specifically at telomeres (Figs 1C and 1D), which can lead to increased DNA damage (Figs 1E and 1F; Step I, Figure S1); (2) The induced telomeric DNA damage then serve as substrates for the formation of dicentric chromosomes (Figs 3A and 3B; Step II, Figure S1); (3) When the dicentric chromosomes remain unresolved and persist into anaphase, they manifest as anaphase bridges (Figs 3C and 3D; Step III, Figure S1); (4) The unresolved anaphase bridges subsequently lead to prolonged mitosis and cytokinesis and the formation of ITCBs (Fig 4; Step IV, Figure S1). Taken together, our data firmly establishes that the dCas9/sgTelo system can be used as a novel cellular model for the BFB cycle.

### cNHEJ, HDR, and BIR promote the formation of ITCBs in the dCas9/sgTelo cells, while TMEJ inhibits them

Given the consistent detection of approximately 3% of the dCas9/sgTelo cells forming ITCBs (Figs 4A and 4B), we then used the ITCB formation as a functional readout of the BFB cycle. We investigated the role of different DSB repair pathways in their formation. Depending on the cell cycle stage and the availability of a source of homology, DSBs in eukaryotes can be repaired by either NHEJ or homology-dependent repair (HDR) ^36^. NHEJ consists of at least two subtypes, cNHEJ, which is dependent on Ku70/80, DNA-PKcs, and Ligase IV, and TMEJ, which is dependent on Pol θ, XRCC1, and Ligase III ^22^. HDR also consists of a few sub-types, including double Holiday Junction (dHJ), synthesis-dependent strand-annealing (SDSA), and break-induced replication (BIR). Using either siRNA (Figure S17) or small molecule inhibitors, none of which lead to pronounced impacts on the cell cycle profiles (Figure S18), we demonstrated that inactivation of proteins involved in HDR (Rad51, Rad52, BRCA1, and BRCA2; Figs 6A to 6C), BIR (POLD3, PIF1 and BLM; Figs 6D and 6E), or cNHEJ (DNA-PKcs; Fig 6F) attenuated the formation of ITCBs. Meanwhile, the inactivation of Polθ (Fig 6G) promoted their formation. The corollary to this finding is that the normal presence of Polθ suppresses the formation of ITCBs, which is somewhat surprising. We wondered if inhibition of Polθ could have shifted the onus of DNA repair to the other available DSB repair pathways. Therefore, we further examined the activity of HDR and cNHEJ in the Polθi treated dCas9/sgTelo cells. Intriguingly, we found that inhibition of Polθ led to increased Rad51 foci and decreased phospho-DNA-PKcs (pDNA-PK) foci and 53BP1 foci (Fig 7). These results suggest that when TMEJ is blocked, telomeric breaks are preferentially repaired through HDR rather than cNHEJ to promote the formation of dicentric chromosomes, anaphase bridges, and ITCBs.

**Figure 6.**
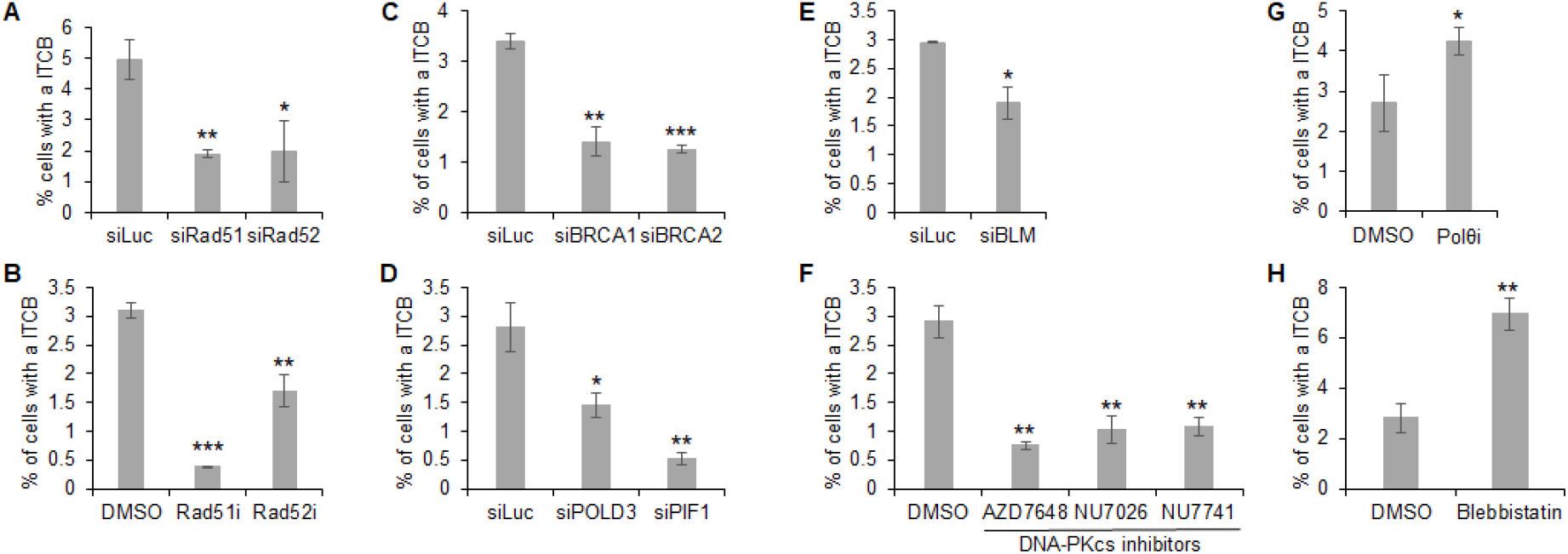
Multiple DNA double-stranded break (DSB) repair pathways regulate the formation of intercellular telomeric chromosome bridges. (**A, C, D, and E**) dCas9/sgTelo cells were transfected with different siRNA twice. 24 hours after the second transfection, cells were treated with doxycycline and Shield1 for 21 hours before they were fixed and imaged. (**B, F, G, and H**) dCas9/sgTelo cells were pre-treated with doxycycline and Shield1 for 6 hours. Small molecule inhibitors targeting different proteins were then added to the growth medium. Cells were grown for another 15 hours before they were fixed and imaged. More than 3000 cells were counted. Error bars are standard deviations from three independent experiments. Standard two-tailed Student’s t-test: ^*^: p < 0.05, ^**^: p < 0.01, ^***^: p < 0.005.

**Figure 7.**
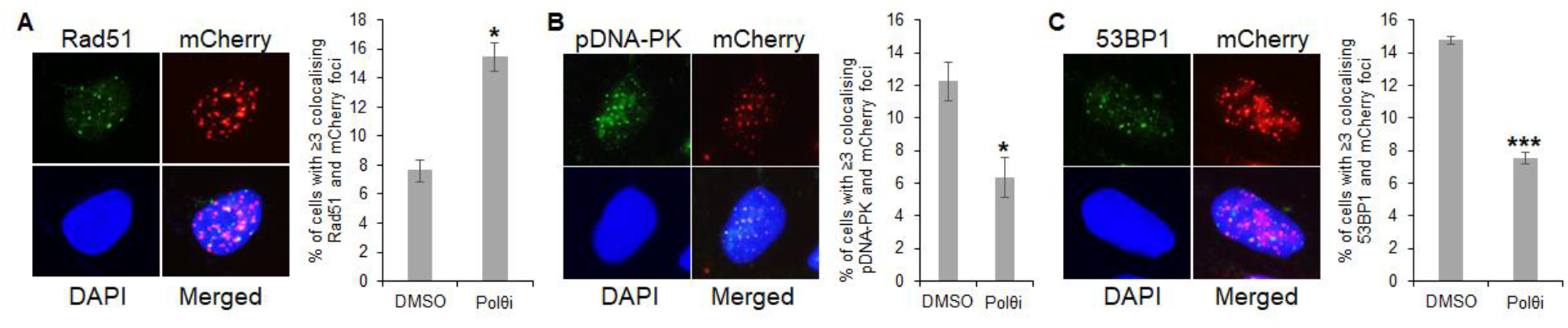
Inhibition of Polθ leads to increased HDR activity and decreased NHEJ activity. dCas9/sgTelo cells were pre-treated with doxycycline and Shield1 for 6 hours. Polθ inhibitor was then added to the growth medium. Cells were grown for another 15 hours before they were fixed and stained with antibodies recognizing either Rad51 (**A**), phosphor-DNA-PKcs (**B**, pDNA-PK), or 53BP1 (**C**). More than 300 cells were analyzed. Error bars are standard deviations from two independent experiments. Standard two-tailed Student’s t-test: ^*^: p < 0.05. ^**^: p < 0.01. ^***^: p < 0.005

Finally, we investigated the mechanism of the breakage of ITCBs. A recent study by Pellman and colleagues suggests that the intercellular chromosome bridge is primarily broken via forces utilizing the actomyosin-dependent mechanical stretching^13^. Consistent with their findings, inhibition of myosin II led to a 3-fold increase of ITCBs (Fig 6H), indicating that many of the ITCBs in the dCas9/sgTelo system are also broken off by the actomyosin-dependent mechanical stretching.

Collectively, our data demonstrate that multiple DSB repair pathways are implicated in forming dicentric chromosomes, anaphase bridges, and ITCBs. cNHEJ and HDR/BIR promote the formation of ITCBs, while the TMEJ suppresses their formation.

## Discussion

As predicted by McClintocks’ original studies, chromosomes experiencing telomeric loss can undergo BFB cycle initiation. However, the precise conditions that create a cellular environment which enables telomere fusion are still unknown. Given that the BFB cycle has such significant implications for tumorigenesis and tumor evolution, understanding it in greater detail is of the utmost importance and this requires additional cellular models ^4, 37^. In this study, we established the dCas9/sgTelo system as an additional cellular model for the BFB cycle, which will complement nicely the current BFB cycle models. First, we demonstrated that targeting dCas9 to telomeres impedes DNA replication and induces pronounced and localized replication stress. This replication stress then induces DNA damage and DSBs at telomeres (Step I of the BFB cycle, Figure S1), which subsequently lead to the formation of dicentric chromosomes (Step II of the BFB cycle, Figure S1), anaphase bridges (Step III of the BFB cycle, Figure S1), and intercellular chromosome bridges (Step IV of the BFB cycle, Figure S1). In addition, we also demonstrated that breakage of the ITCBs in the dCas9/sgTelo system is partially due to the actomyosin-dependent mechanical stretching, consistent with the previous finding from Pellman’s group ^13^. Using this newly established BFB cycle model, we showed that multiple DSB repair pathways are implicated in the formation or suppression of dicentric chromosomes, anaphase bridges, and the ITCBs. Specifically, we found that cNHEJ, HDR, and BIR promote the formation of ITCBs while TMEJ suppresses them. Finally, utilizing various molecular and cellular assays, we uncovered many novel features of the intercellular chromosome bridges. These include discovering the DNA knots in the middle of many ITCBs and the active DNA synthesis/repair at many DNA knots. The findings presented here will profoundly impact the understanding of how the BFB cycle affects tumorigenesis, tumor evolution, and drug resistance.

### dCas9/sgRNA as a versatile and locus-specific tool for investigating replication stress in vivo

DNA damage can be induced by various endogenous or exogenous sources ^36, 38^. Replication stress is one of the most common endogenous sources of DNA damage, especially at those difficult-to-replicate (DTR) genomic loci, including centromeres, common fragile sites, and telomeres, which contribute to the development of many human diseases including cancer ^39-41^. The source of the replication stress at these sites can include factors such as dysregulated DNA repair, nucleotide imbalances, fork barriers etc. Replication fork barriers include both exogenous and endogenous sources and are of particular interest in the context of this study. Exogenous sources include exposure to UV radiation, or compounds such as cross-linking agents (e.g., mitomycin C and cisplatin), or topoisomerase inhibitors (i.e., camptothecin) and others. Endogenous fork barriers can include secondary DNA:RNA hybrid structures or DNA structures like G-4 quadraplexes.

Much of the research done on sources of replication stress and cellular response to replication stress relies on the exogenous replication stress inducers (RSI). The disadvantages of using these exogenous RSIs include (1) inducing DNA damage via non-DNA replication-related mechanisms and (2) indiscriminately blocking DNA replication at many genomic regions/loci. This in turn has limited the understanding of effects of specific localized replication stress responses, making new replication stress models of utmost importance. To address these limitations, a few novel endogenous and locus-specific replication fork blocks (RFBs) via strong protein-DNA interactions, including LacO-LacI, TetO-TetR, and Tus-Ter, have been developed in mammalian cells ^42-44^. In addition, we recently demonstrated that depletion of FANCM in cells that adopted the ALT pathway induces a pronounced replication stress response at telomeres, and we thus proposed that the FANCM deficiency-induced replication <u>s</u>tress at <u>A</u>LT telomeres (MR-SAT) can be used as a telomere-specific DNA replication perturbation model ^29, 45^. Using the MR-SAT system, we uncovered a novel DNA damage checkpoint function of POLDIP3 ^46^. Here, we demonstrated that targeting dCas9 to telomeres via sgTelo also induced a robust replication stress response at telomeres (Fig 1C), suggesting that the dCas9/sgRNA system is potentially the most versatile tool to induce replication stress at any genomic locus and both in vitro ^28^ and in vivo (this study). In addition, as demonstrated here, the dCas9/sgRNA system can also be used to investigate the downstream events of the replication stress response. For example, the formation of dicentric chromosomes, anaphase bridges, intercellular chromosome bridges, and potential genomic changes in the two daughter cells (Steps VI and VII in Figure S1) and their progenies.

### Not all telomeres behave the same: chromosome arm-specific response to replication stress

One of the intriguing findings in this study is the differential responses to replication stress at telomeres among different chromosome arms and even between the two arms of the same chromosome (Fig 2, Figs S6 to S10, and Table 1). Given the 21-hour induction time, it is not entirely surprising that at the population level, some of the characteristics studied exhibited no statistically significant differences between dCas9/sgNS and dCas9/sgTelo cells. On the other hand, we did notice the stark differences on individual chromosome arms for certain telomere/chromosome end features. For example, though there is no significant difference in the mean length of End Tel between the dCas9/sgNS cells and the dCas9/sgTelo cells, the telomeres of chromosome arms 18p and 20q reduce significantly in the dCas9/sgTelo cells compared to the dCas9/sgNS cells (Figure S6). Conversely, the telomere of chromosome arm 14q is longer in the dCas9/sgTelo cells compared to the dCas9/sgNS cells (Figure S6). One potential mechanism that regulates the chromosome arm-specific replication stress response may be differential expression of <u>te</u>lomeric <u>r</u>epeat-containing <u>R</u>N<u>A</u> (TERRA) or the formation of different amounts of TERRA R-loops. For example, several groups showed that different chromosome arms express different amounts of TERRA ^47-53^. We and others have demonstrated that aberrant accumulation of TERRA R-loops at telomeres can induce a robust replication stress response in ALT cells ^54-55^. Other possible reasons include the unique epigenetic and chromatin environments, the three-dimensional location within the nucleus, or the interaction/regulation of chromosomes with nuclear peripheral proteins. We do want to note that to define all the chromosome arm specific features with high confidence, including TFE, fusion/ITS-, and fusion/ITS+, we will need to collect many more molecules in the future.

### Initiation of the BFB cycle by the replication fork stalling-/collapse-induced DSBs

One of the unanswered questions related to the BFB cycle is: how the initial DSB(s) in Step I of the BFB cycle is induced (Figure S1)? It has been well recognized that the DTR loci, including telomeres, are more prone to DNA damage than other regions of the genome ^39-41^. The primary cause of DNA damage at these DTR loci is likely due to the frequent replication fork pausing, stalling, or even collapsing, often due to the inherent sequences of the regions which can be prone to forming secondary structures. This in turn leads to the activation of the replication stress response and DNA damage. The data presented here indicates that telomeric replication stress-induced DSBs are likely the primary endogenous source of DSBs to initiate the first BFB cycle. Furthermore, we showed that cNHEJ, HDR, and BIR promote the formation of ITCBs (Fig 6). BIR has previously been shown to be activated by a single one-ended DSB ^56^. Therefore, in theory, one unrepaired DSB from a single telomere has the latent potential to trigger BFB cycle with far reaching genomic consequences.

Finally, we demonstrated that inactivation of the TMEJ pathway led to an increase in the formation of ITCBs (Fig 6G). Consistent with our observation, two previous studies observed an increase in the frequency of inter-chromosomal translocation in Polθ deficient cells either induced by CRISPR/Cas9 ^57^ or at the endogenous Myc/IgH loci ^58^. In addition, Davis and colleagues demonstrated that Polθ deficient cells manifested an increase in the inter-homolog recombination (IHR), often leading to loss-of-heterozygosity ^59^. Collectively, these data suggest that TMEJ may play an essential role in suppressing tumorigenesis and tumor evolution by preventing the formation of dicentric chromosomes, thus the initiation of the BFB cycle through inhibiting IHR and inter-chromosomal translocation (Step II in the Figure S1).

### Novel molecular and structural features of the intercellular chromosome bridges

Detection of an intercellular chromosome bridge in the intercellular canal during cytokinesis activates the abscission checkpoint to allow the cell more time to resolve the bridge ^60^. Because of the rarity of the intercellular chromosome bridge, its molecular and structural features remain largely unknown. Utilizing the dCas9/sgTelo-induced ITCBs, we have uncovered several novel features of the intercellular chromosome bridge. First, we discovered DNA knots in the middle of 86% of the ITCBs, which also colocalize with the midbody (Fig 4A). Most intriguingly, we demonstrated that approximately one third of the DNA knots are sites of active DNA repair and/or synthesis as indicated by EdU incorporation (Fig 4A-iv and Figure S12). While this study was ongoing, the Zachos’ group reported their discovery of the DNA knots on either spontaneously generated or HU-induced intercellular chromosome bridges ^61^. Consistent with their findings, we also detected Top2A at the DNA knots (Fig 4A-v and Figure S13), suggesting that over-twisted and/or catenated DNA is likely present in many ITCBs. In addition to Top2A, we also detected many other DDR proteins at different regions of the ITCBs, including the DNA knots (Fig 4 and Figure S14). For example, the ssDNA binding protein, RPA, was detected at both the bodies and DNA knots of the ITCBs, suggesting that there are ssDNA gaps at both places. In addition to RPA, we also detected 53BP1, FANCI, and cGAS on the body of ITCBs. Interestingly, a previous study from Mitchison’s group also detected cGAS on the body of the intercellular chromosome bridges ^62^. Their studies suggest that the intercellular chromosome bridge facilitates the activation of cGAS after drug-induced mitotic errors in human cells. The only protein that can be detected throughout the ITCBs is 53BP1. Its precise role in the formation and resolution of the ITCBs is currently under intense investigation.

In summary, the discoveries presented here have filled many knowledge gaps in the understanding of the BFB cycle. Because of the crucial role of the BFB cycle in the continued tumor evolution ^4-5^, which could lead to more significant tumor heterogeneity, treatment resistance, and relapse, our discoveries could also have significant implications in cancer treatments. For example, by suppressing the formation of ITCBs, cNHEJ, HDR, and BIR inhibitors may slow down continued tumor evolution, reducing treatment resistance and tumor relapse. Conversely, since Polθ inhibitor promotes the formation of ITCBs, it may accelerate tumor evolution, leading to increased treatment resistance and accelerated tumor relapse.

## Acknowledgments

We want to thank Dr. Erik Sontheimer for generously providing both U2OS-dCas9/sgNS and U2OS-dCas9/sgTel cells. We also want to thank Dr. James Haber for reading our manuscript and providing insightful comments. Time-lapse movies were acquired in the AIF at Albert Einstein College of Medicine by Dr. Vera DesMarais using the AIF DVCore microscope. The AIF is partially funded by the NCI Cancer Center support grant P30CA013330.

## Data Availability

raw and processed data reported in this paper will be available upon request.

## Code Availability

N/A.

## Author contributions

DZ conceived the idea. Investigation: MS, KR, AMP, KL, SD, NB, NG, HZA, LUB; Data analysis: DZ, MX, MS, KR, AMP, KL, SD, NB, NG, HZA, LUB, RFS; CS provided advice and inputs. All of the authors reviewed, edited, and approved the manuscript.

