## Supplemental Information for "Elucidation of the molecular mechanism of the breakage-fusion-bridge (BFB) cycle using a CRISPR-dCas9 cellular model"

**Supplementary Information (SI)**

**Supplemental Figure Legends**

**Figure S1**. **A model of the Breakage-Fusion-Bridge (BFB) cycle**. Step I: generation of one or more double-stranded DNA breaks (DSBs) as marked by a flash. Step II: two chromosome arms, either from a pair of sister chromatins (top) or two non-sister chromosomes (bottom), are fused together by one or more DSB repair pathways. Step III: anaphase bridge formation from a dicentric chromosome with two functional centromeres. Step IV: formation of an intercellular chromosome bridge during cytokinesis from one or more unresolved anaphase bridge. Step V: breakage of the intercellular chromosome bridge. Step VIa: initiation of another round of the BFB cycle by forming one or more dicentric chromosomes. Steps VIb and VII: healing of the broken chromosome via chromothripsis.

**Figure S2**: **Additional DNA damage response proteins are recruited to the damaged telomeres in the dCas9/sgTelo cells**. 21 hours after the addition of Dox and Shield1, dCas9/sgTelo cells were fixed and stained with either a PNA probe recognizing telomeres (TelC) (A), or different antibodies recognizing TRF1 (**A**) or as indicated (**B** to **G**).

**Figure S3**: A diagram of the three-colored SMTA-OM (**A**) and its readouts (**B**).

**Figure S4**: All the DNA molecules detected by the three-color SMTA-OM and assigned to the chromosome arm 3q from both dCas9/sgNS and dCas9/sgTelo cells.

**Figure S5**: All the DNA molecules detected by the three-color SMTA-OM and assigned to the chromosome arm 8q from both dCas9/sgNS and dCas9/sgTelo cells.

**Figure S6**: **Analysis of mean length of End Tel**. End Tel accounts for 72.8% of the molecules in the dCas9/sgTelo cells and 89.1% of the molecules in the dCas9/sgNS cells (Supplementary tables 1 and 2). We calculated the mean length of End Tel for each chromosome arm. The overall mean length of End Tel is 11.3 kb for the dCas9/sgTelo cells and 11.7 kb for the dCas9/sgNS cells (**A** and Supplementary tables 1 and 2). The two-tailed paired t-test results in a *p* value of 0.248, suggesting that there is no significant difference in the overall mean length of End Tel between the two samples. To examine whether there is any chromosome arm-specific changes, we produced a scatter plot, in which the differences in mean length of End Tel (sgTelo-sgNS) are plotted against the mean length of End Tel ratios (sgTelo/sgNS) for each chromosome arm (**B**). Overall, there were equal numbers of chromosome arms showing an increase, i.e., the sgTelo/sgNS ratio is above 1 and the difference (sgTelo-sgNS) is above 0 and those showing a decrease, i.e., the sgTelo/sgNS ratio is below 1 and the difference (sgTelo-sgNS) is below 0. (**C**) Individual chromosome arms manifested significant changes in the mean length of End Tel. 18p and 20q have an identical ratio of 0.81 meanwhile have a difference of 2.6 and 2.7, respectively, which indicate that there is a decrease in the mean length of End Tel in the dCas9/sgTelo cells compared to that in the dCas9/sgNS cells. 14q has a slight increase in the mean length of End Tel in the dCas9/sgTelo cells, with a ratio of 1.16 and a difference of 1.83 compared to that in the dCas9/sgNS cells. To confirm the three outlier arms, we performed a t-test using the end telomere length measurements for each arm from the dCas9/sgTelo cells against the corresponding measurements of the same arm from the dCas9/sgNS cells. With a high confidence *p-value* ($p < 0.05$), these outlier arms are confirmed to have statistically significant differences: 14q ($p$=0.047), 18p ($p$=0.010), and 20q ($p$=0.005).

**Figure S7**: **Analysis of telomere free ends, or TFEs**. dCas9/sgTelo cells manifest an elevated percentage of TFEs compared to the dCas9/sgNS cells (Supplementary tables 1 and 2). Approximately 8% of the molecules in the dCas9/sgTelo cells and 5% of the molecules in the dCas9/sgNS cells are TFEs. A two-tailed paired t-test was performed and resulted in a P value of 0.04.

**Figure S8**: **Analysis of fusion/ITS+ (ITS+)**. ITS+ is highly elevated in the dCas9/sgTelo cells compared to the dCas9/sgNS cells (Supplementary tables 1 and 2). ITS+ was detected in 15.0% of the molecules in the dCas9/sgTelo cells compared to 5.5% of the molecules in the dCas9/sgNS cells (**A**). A two-tailed paired t-test was performed using the ITS+ percentages of all the chromosome arms in the dCas9/sgTelo cells and the dCas9/sgNS cells, resulting in a $P$ value of 0.0001. To further examine which arm experiences a significant change of ITS+, we generated a scatter plot using the differences in ITS+ percentage between dCas9/sgTelo and dCas9/sgNS for each chromosome arm (sgTelo ITS+% - sgNS ITS+%) against the ratio of the percentages of each chromosome arm of the two samples (sgTelo ITS+% / sgNS ITS+%) (**B**). For the arms that show little to no differences in ITS+, the data points fall near (0,1). For the arms that show a higher percentage of ITS+ in the dCas9/sgTelo cells than that of the dCas9/sgNS cells, the ratio is above 1, while the difference is bigger than 0. The blue dots indicate that ITS+ is found only in the dCas9/sgTelo cells, in which the ratio would be infinity. Instead, we used the number "12" to represent the infinity ratio. The red dot represents that ITS+ is found only in the dCas9/sgNS cells; thus, the ratio would be 0. Most chromosome arms manifest the ratio above 1 and the difference above 0, indicating that ITS+ is elevated in the dCas9/sgTelo cells compared to the dCas9/sgNS cells. (**C**) 8q showed a dramatic increase in the dCas9/sgTelo cells comparing to the dCas9/sgNS cells.

**Figure S9**: **Analysis of fusion/ITS- (ITS-)**. Overall, ITS- are relatively lower abundance events than ITS+. 4.0% of the molecules in the dCas9/sgTelo cells and 0.4% of the molecules in the dCas9/sgNS cells were identified as ITS- (**A**, Supplementary tables 1 and 2). A two-tailed paired t-test results in a P value of 0.079, which is likely due to the fact that only a few chromosome arms in the dCas9/sgNS cells have ITS- (Supplementary tables 1 and 2). A scatter plot shows the differences in ITS- among different chromosome arms (**B**). For example, there is no ITS- detected for 3q in the dCas9/sgNS cells while a striking 47.6% of the molecules are ITS- in the dCas9/sgTelo cells (**C**).

**Figure S10**: **Analysis of the mean length of telomeres in the fusion/ITS+**. (**A**) Overall, there is no significant difference in the mean length of telomeres in ITS+ between the dCas9/sgTelo cells and dCas9/sgNS cells (Supplementary tables 1 and 2). A two-tailed paired t-test results in a P value of 0.869. The lengths of telomeres in ITS+ in the dCas9/sgTelo cells range from 4.4 kb to 22.9 kb with an average of 11.8 kb. The telomere lengths in ITS+ in the dCas9/sgNS cells range from 5.4 kb to17.5 kb with an average of 11.2 kb. (**B**) A scatter plot of the mean length of telomere differences (sgTelo-sgNS) vs the mean length of telomere ratios (sgTelo/sgNS) for each chromosome arm. Even though there is no overall differences in the mean length of telomeres in the fusion/ITS+, 8q showed a significant decrease in the dCas9/sgTelo cells compared to that in the dCas9/sgNS cells, while 19p showed a significant increase (**C**).

**Figure S11**: **DNA knots and the staining pattern of 53BP1 on the ITCBs**. (**A**) 21 hours after the addition of Dox and Shield1, dCas9/sgTelo cells were fixed and stained with an antibody recognizing 53BP1. All nuclei were also stained with DAPI. (**B**) 50 ITCBs from (A) were analyzed.

**Fig S12. EdU pulse labeling assay.** (**A**) Zoom out images from 15 min EdU pulse labeling. The white arrow heads mark two ITCBs. (**B**) Two representative ITCB images from 30 min and 15 min EdU pulsing, respectively. The white arrows mark the DNA knots. The orange arrows mark two micronuclei. (**C**) The percentage of EdU positive DNA knots. More than 30 ITCBs were analyzed.

**Figure S13**: **More than half of the DNA knots are positive of TOP2A**. (**A**) 21 hours after the addition of Dox and Shield1, dCas9/sgTelo cells were fixed and stained with an antibody recognizing Top2A. All nuclei were also stained with DAPI. (**B**) More than 50 ITCBs from (A) were analyzed.

**Figure S14**: **Additional DNA damage response proteins are found at the intercellular telomeric chromosome bridges**. (**A** to **M**) 21 hours after the addition of Dox and Shield1, dCas9/sgTelo cells were fixed and stained with antibodies recognizing different proteins as indicated. All nuclei were also stained with DAPI. (**N**) A summary of the staining patterns of different proteins.

**Figure S15**. **Structural features of the intercellular telomeric chromosome bridges obtained via STORM**. (**A**) The number of branches for the 26 bridges. (**B**) The length of the 26 bridges. (**C**) The width of the 26 bridges.

**Figure S16**. **Time-lapse analysis of live dCas9/sgTelo cells**. Asynchronous dCas9/sgTelo cells were pre-treated with doxycycline and Shield1 for 8 hours. The cells were then monitored using a fully automated DVCore microscope (Leica-microsystems). Images were acquired every 30 minutes for 25 hours as indicated by the Tn. Tn' is the enlarged view of Tn. (**A** and **B**) The red arrows indicate a cell expressing a low level of mCherry and manifesting relatively normal cell cycle progression. (**C**) The red arrows indicate a cell that formed the intercellular telomeric chromosome bridge at T34. (**D**) The red arrows indicate a cell that underwent a mitotic catastrophe.

**Figure S17**. **Depletion efficiency of various siRNA**. dCas9/sgTelo cells were transfected twice with different siRNA. Whole-cell lysates were immunoblotted (IB) with different antibodies, as indicated on the right.

**Figure S18**. **Cell cycle analysis of dCas9/sgTelo cells treated with various siRNA or small molecule inhibitors**. (A to H) dCas9/sgTelo cells were transfected twice with different siRNA. 24 hours after the second transfections, cells were treated with doxycycline and Shield1 for 21 hours before they were stained with propidium iodide (PI). (I to Q) dCas9/sgTelo cells were first treated with doxycycline and Shield1 for 6 hours. DMSO, or small molecule inhibitors targeting RAD51 (B02, or RAD51i), RAD52 (D-I03, or RAD52i), DNA-PKcs (AZD7648, NU7441, NU7076), Pol θ (ART558 and Novobiocin), or myosin II (blebblstatin) were then added to the growth medium. Cells were grown for another 15 hours before they were stained with propidium iodide (PI). PI stained cells were analyzed by flow cytometry.

### Extended videos

**Extended video 1:** The time-lapse of a cell that expresses a low level of mCherry undergoing normal mitosis and cytokinesis (marked by a blue arrow) and a cell undergoing mitotic catastrophe (marked by a yellow arrow).

**Extended video 2:** The time-lapse of a cell that expresses a high level of mCherry undergoing the formation of an intercellular telomeric chromosome bridge (marked by a blue arrow).

### Supplemental Tables

**Supplemental Table 1: The raw data of telomere and chromosome end characterization using the three-color SMTA-OM**. In total, we were able to characterize 35 of 46 chromosome arms. On average, we analyzed approximately 30 molecules per chromosome arm. The acrocentric chromosome arms 13p, 14p, 15p, 21p, and 22p cannot be analyzed because no reference genome is currently available. Fewer than five molecules of 4q or 18q could be assigned to their corresponding consensus sequences, which are too few to be analyzed statistically. 9q, 10p, 12p, and 17q have no consensus contigs assembled. Additional notes: The numbers in black font indicate the telomere length measurement of an End Telomeres or End Tel. The length marked as “0” indicates that the DNA molecule has no detectable telomere, which is thus considered a telomere-free end, or TFE. The numbers in red are telomere length measurements for the fusion/ITS+, or ITS+. The fusion molecule measured as “0” indicates that it has no detectable telomere, i.e., fusion/ITS-, or ITS-.

**Supplemental Table 2: Summary of the characterization of 35 chromosome arms using the three-color SMTA-OM**. We calculated and obtained multiple parameters of 35 chromosome arms, which include (1) the total number of single molecules for each chromosome arm; (2) the mean lengths of telomeres for the End Telomeres and Fusions/ITS+; (3) the standard deviations; (4) the frequencies of end telomere, TFE, fusion/ITS+ (or ITS+), and fusion/ITS- (or ITS-) events.

**Supplemental Table 3:** Structure features of the intercellular telomeric chromosome bridges (ITCBs) obtained via STORM.

**Supplemental Table 4:** Antibodies used in this study.

**Supplemental Table 5:** siRNA used in this study.

**Supplemental Table 6:** Small molecule inhibitors used in this study

**Fig S1:** The BFB cycle

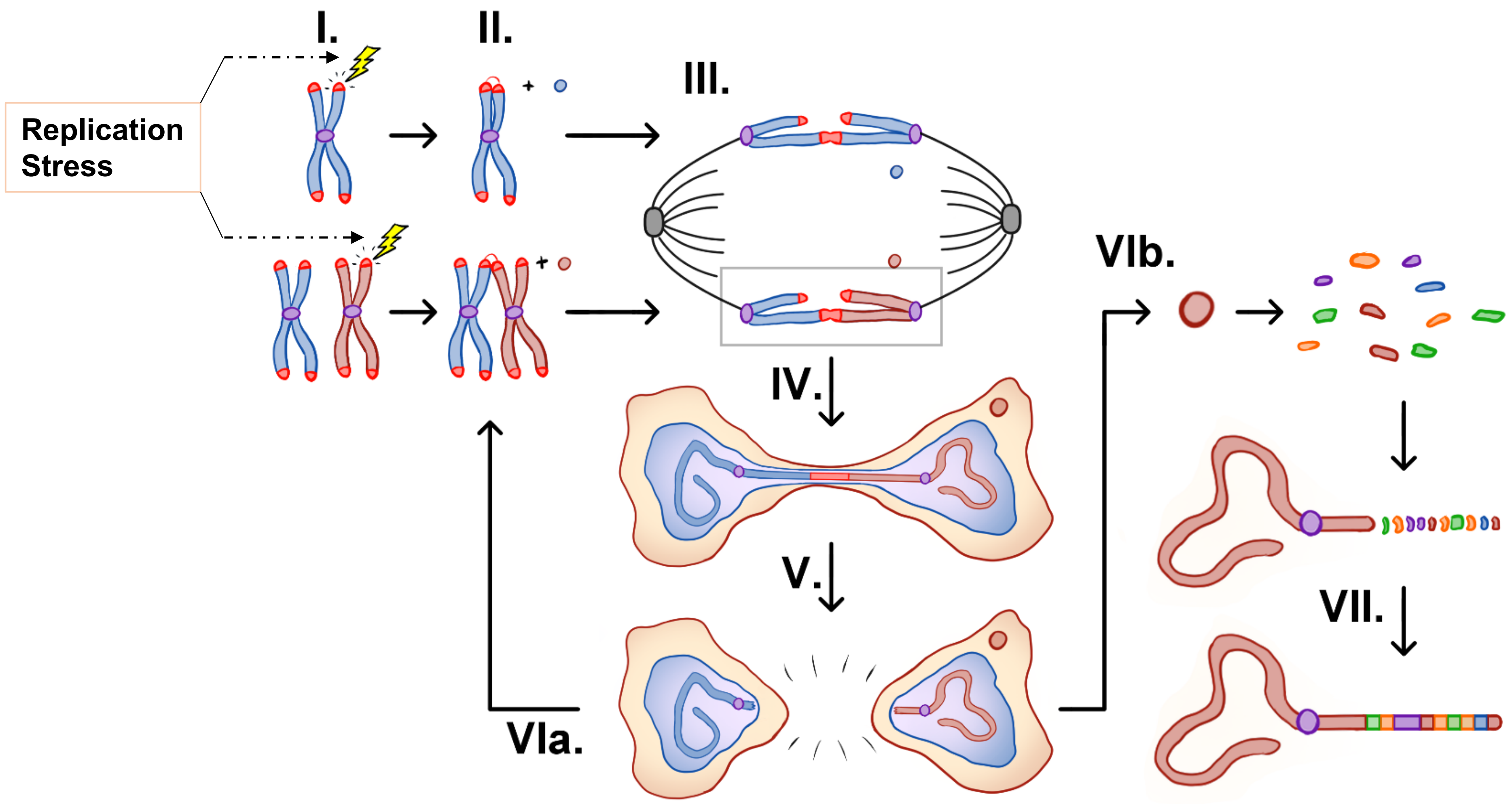

**Fig S2:** Additional DNA damage response proteins are recruited to the damaged telomeres in the dCas9/sgTelo cells.

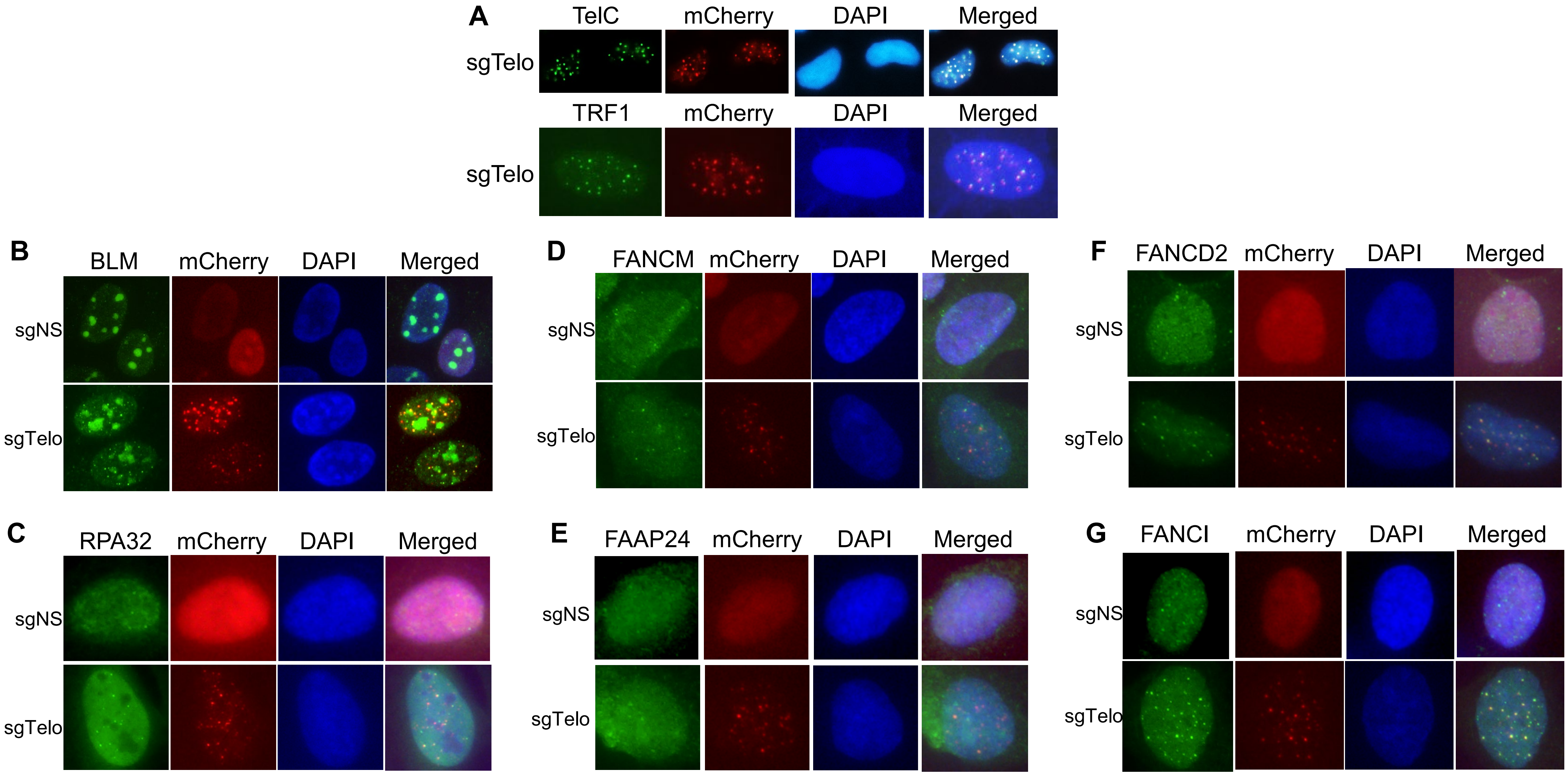

**Fig S3**: A diagram of single molecule telomere assay via optical mapping (SMTA-OM)

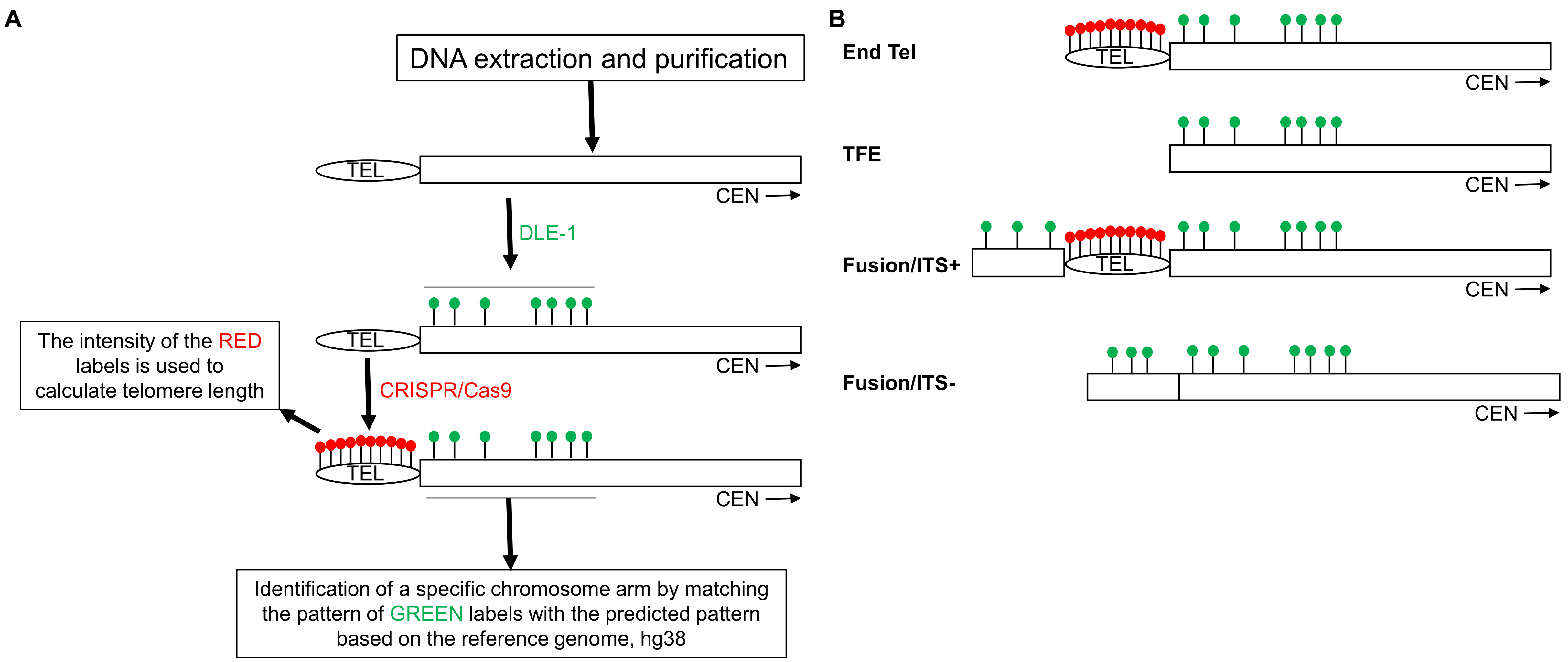

**Fig S4:** All the DNA molecules detected by the three-color SMTA-OM and assigned to chromosome arm 3q from both dCas9/sgNS and dCas9/sgTelo cells.

**A. 3q of the dCas9/sgNS**

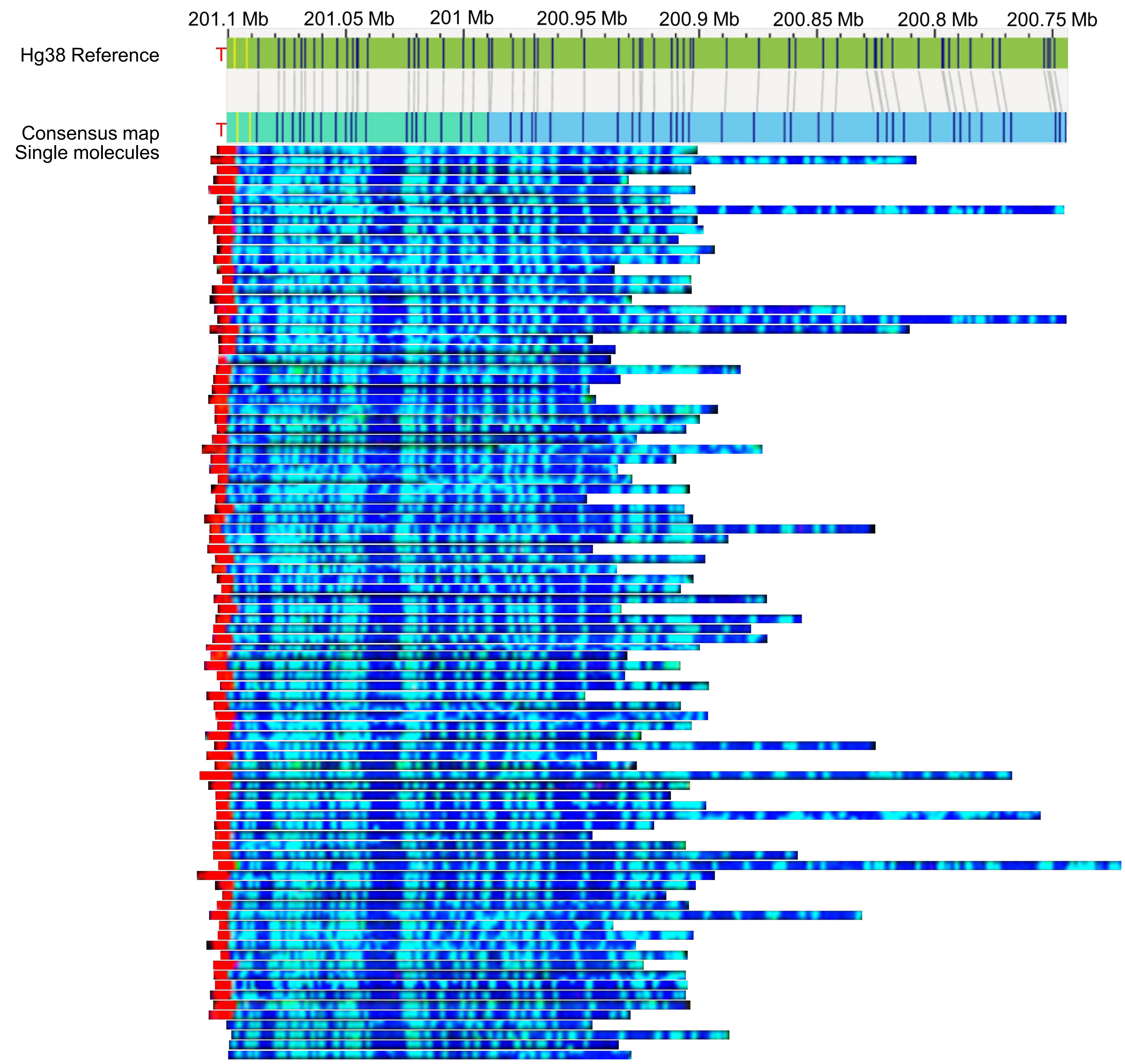


**B. 3q of the dCas9/sgTelo**

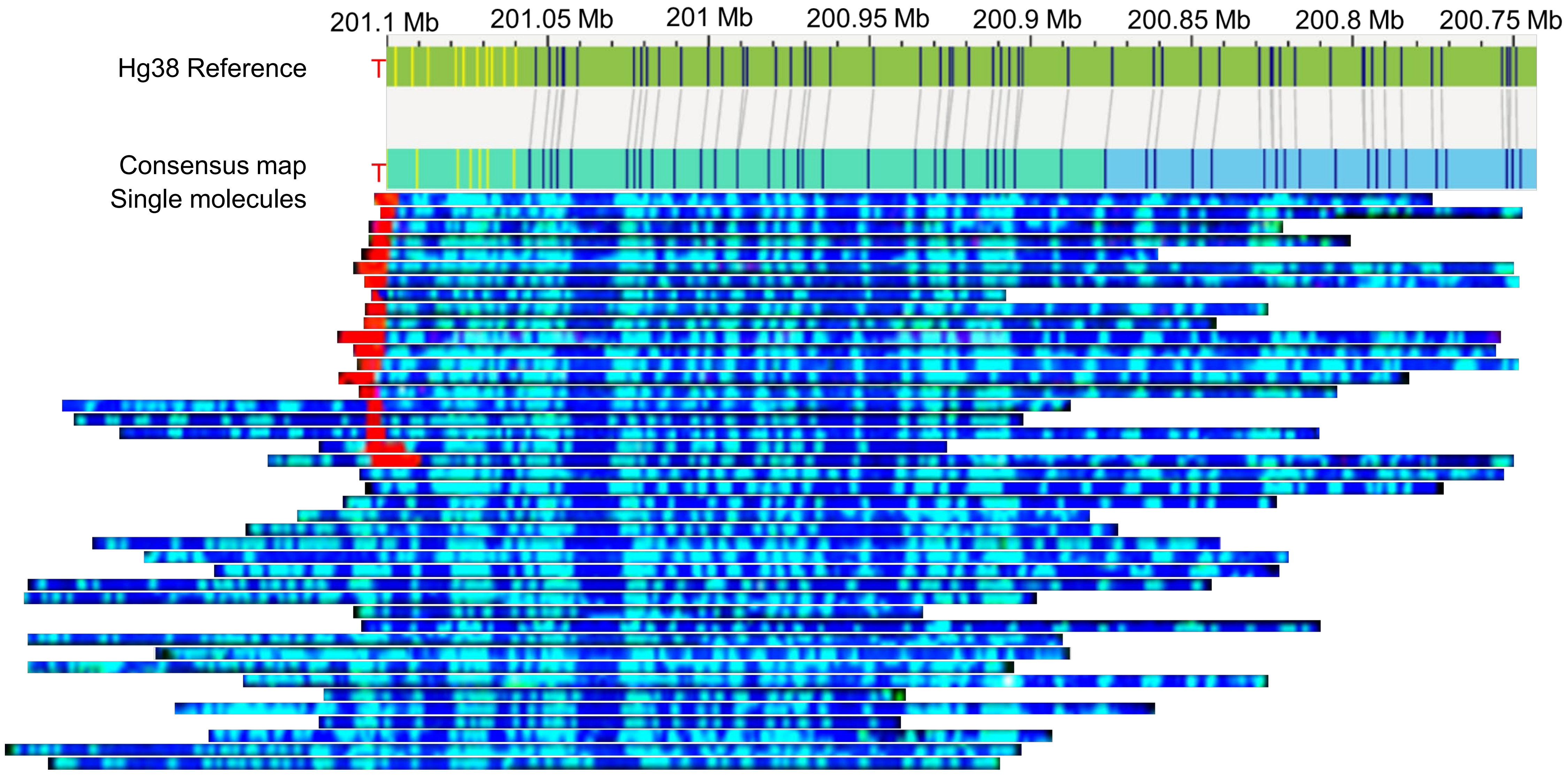

**Fig S5** All the DNA molecules detected by the three-color SMTA-OM and assigned to chromosome arm 8q from both dCas9/sgNS and dCas9/sgTelo cells.

**A. 8q of the dCas9/sgNS**

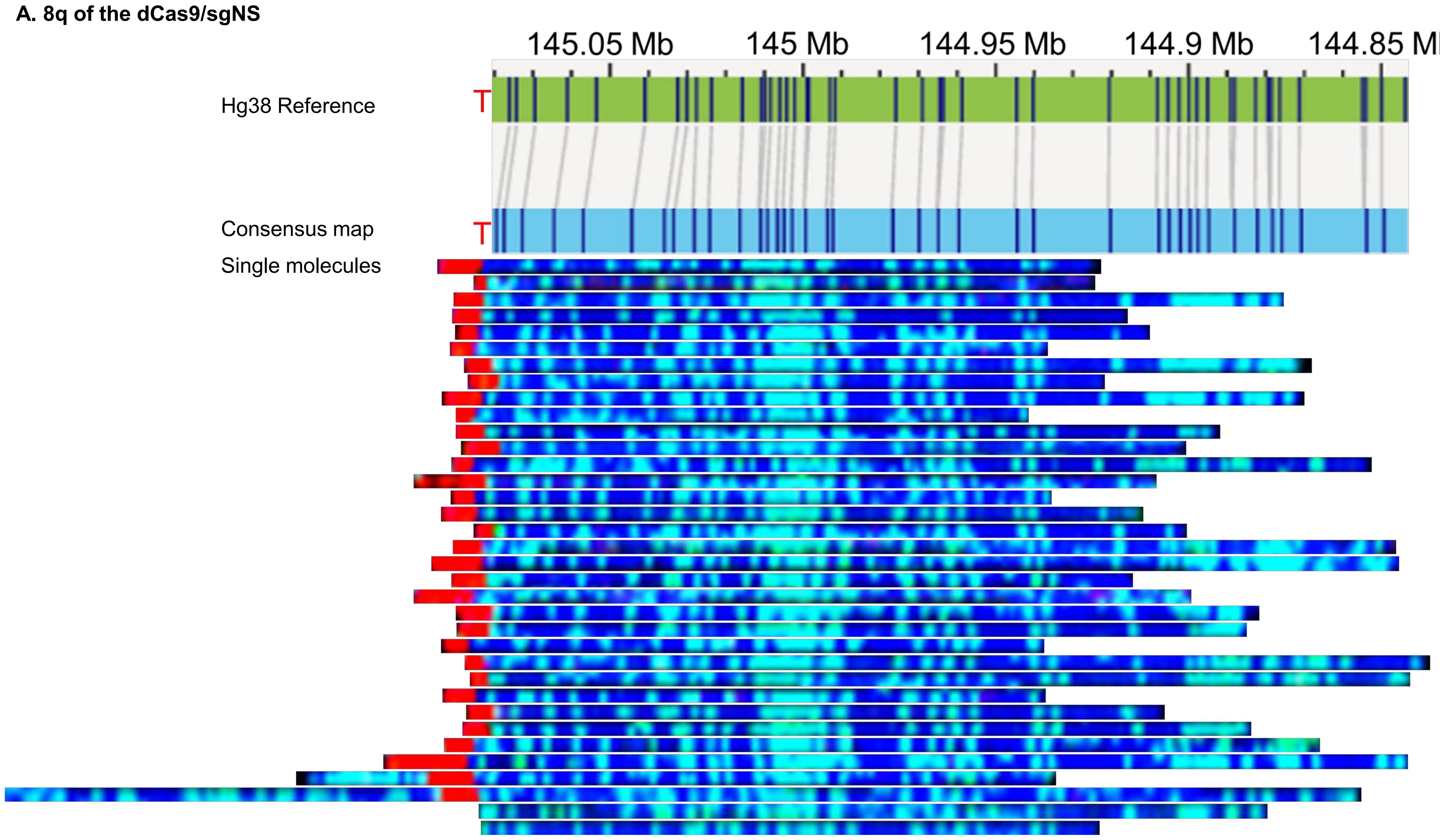


**B. 8q of the dCas9/sgTelo**

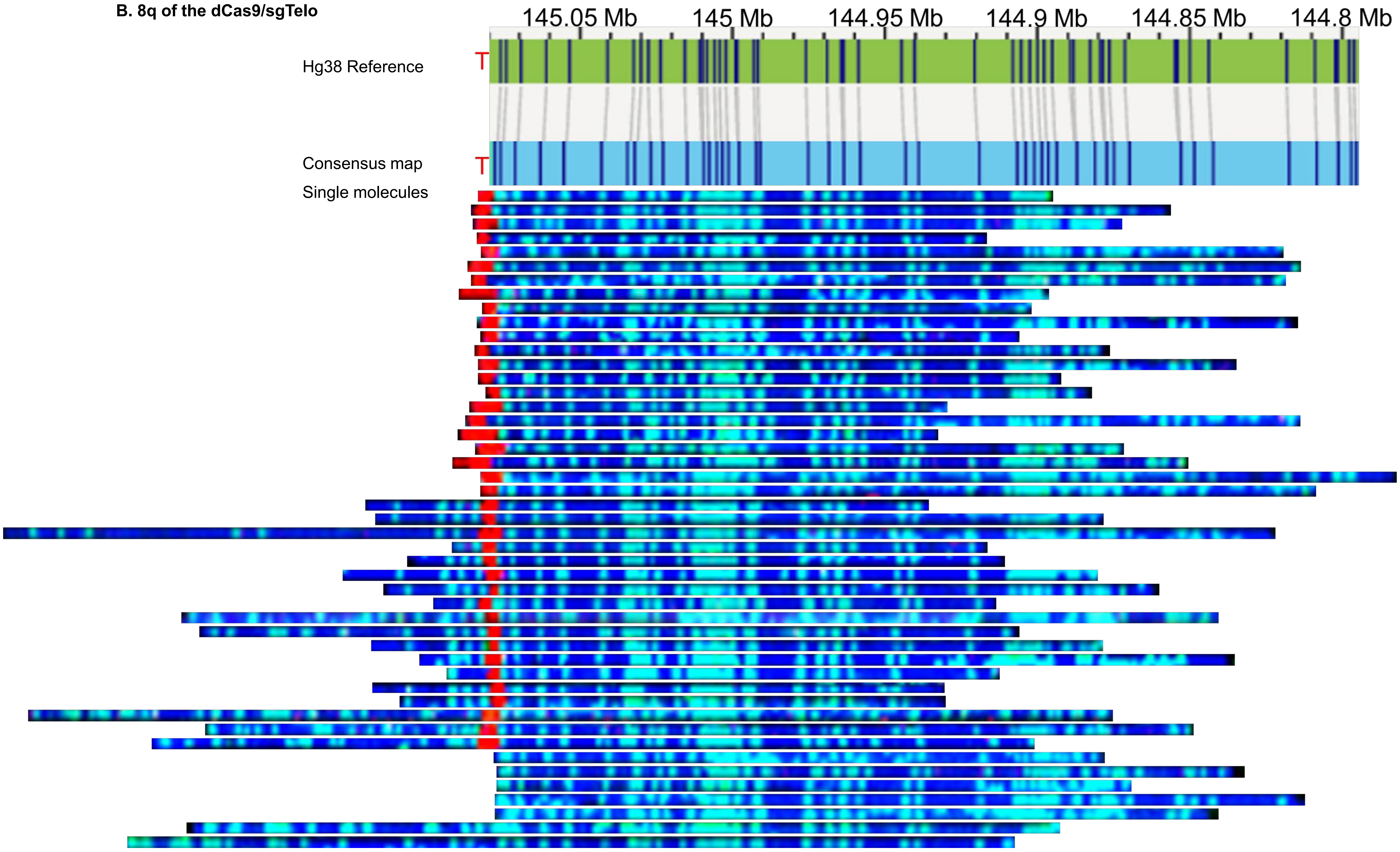

**Fig S6:** Analysis of the mean length of End Tel

**A**

| | Mean length of End Tel | P value |
|---|---|---|
| sgTelo (kb) | 11.3 | 0.248 |
| sgNS (kb) | 11.7 | |

**B**

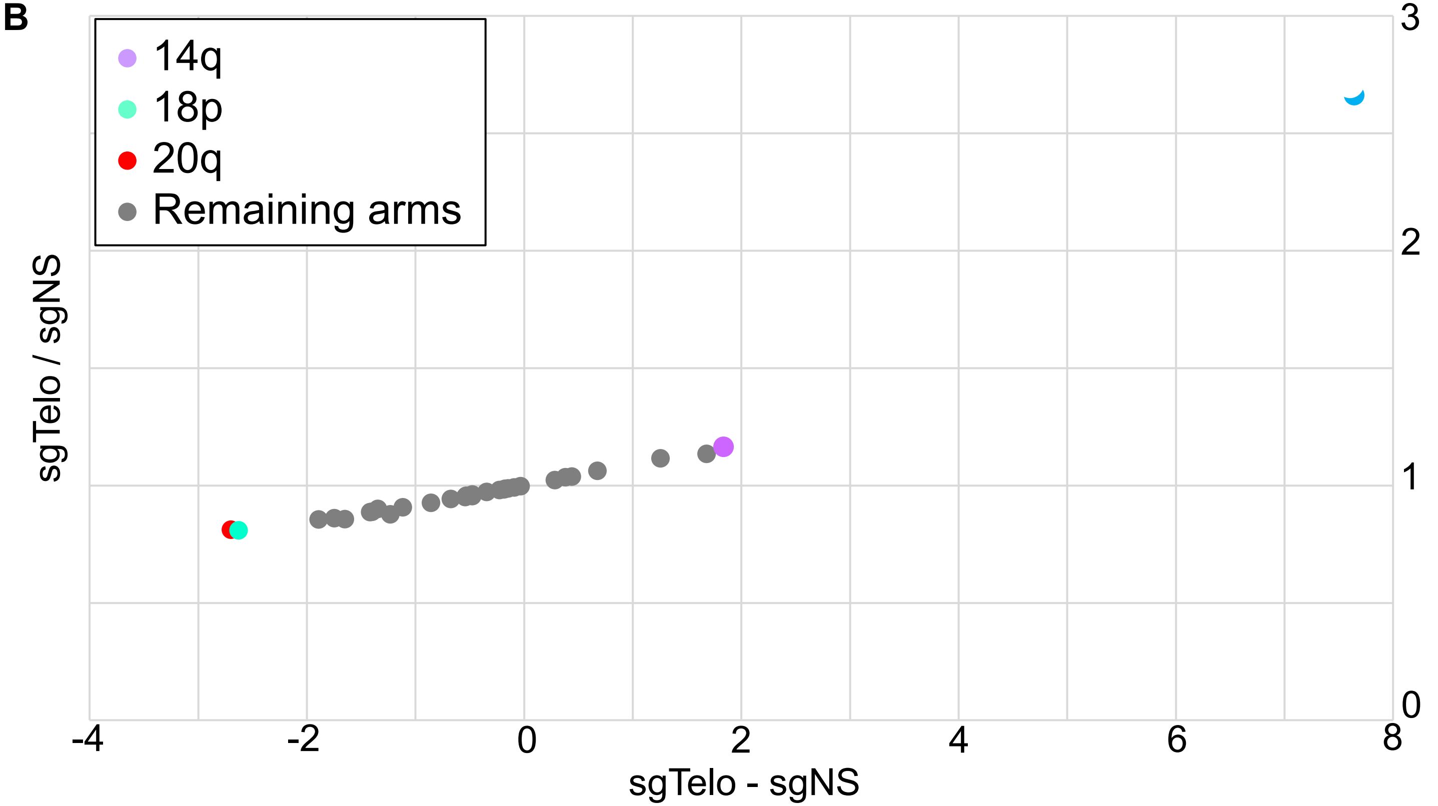


**C**

| Chromosome arm | 14q | 18p | 20q |
|---|---|---|---|
| sgTelo (kb) | 13.0 | 11.2 | 11.6 |
| sgNS (kb) | 11.1 | 13.9 | 14.2 |
| P value | 0.047 | 0.010 | 0.005 |

**Fig S7:** Analysis of the telomere-free ends (TFEs).

**A**

| | TFE % | P value |
|---|---|---|
| sgTelo | 8.0 | 0.0400 |
| sgNS | 5.0 | |

**B**

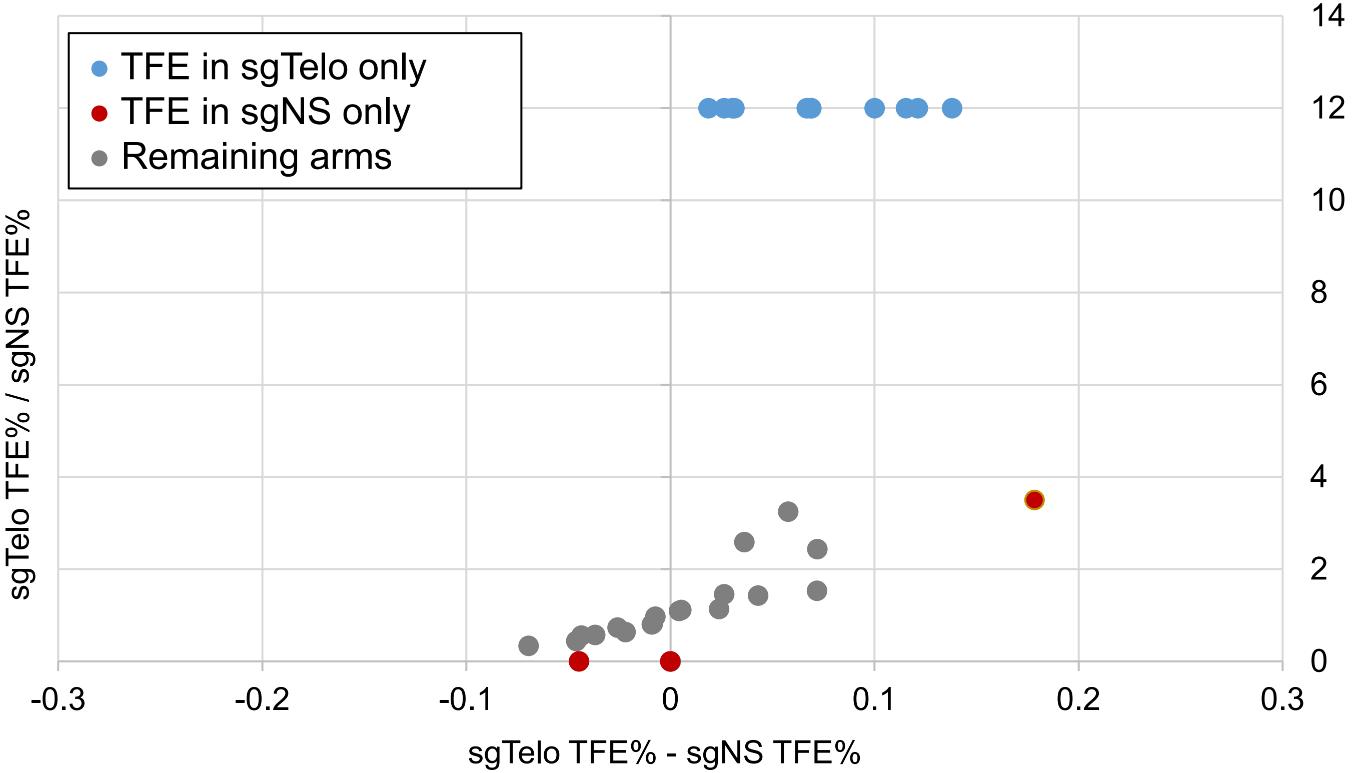

**Fig S8:** Analysis of the fusions/ITS+ (ITS+).

A

| | ITS+ % | P value |
|---|---|---|
| sgTelo | 15.2 | 0.0001 |
| sgNS | 5.4 | |

B

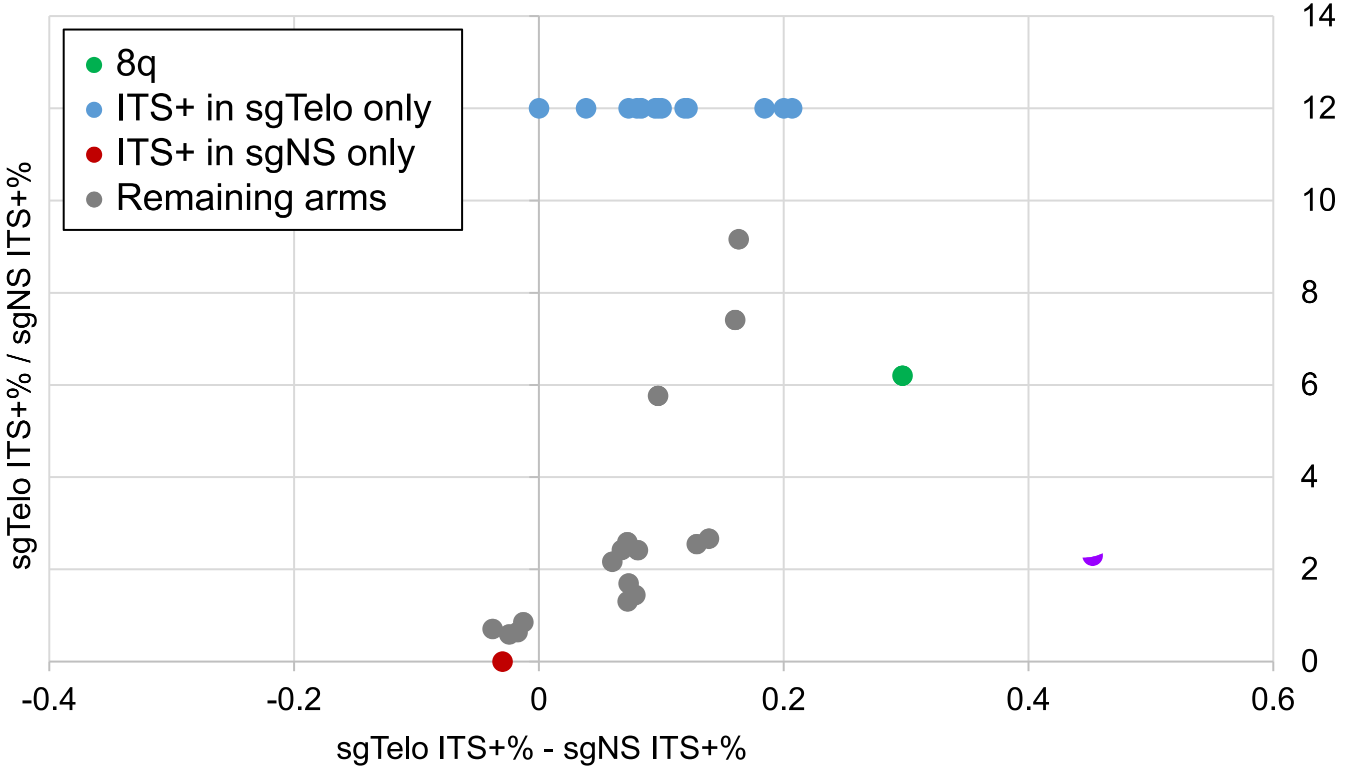


C

| Chromosome arm | 8q |
|---|---|
| sgTelo ITS+ % | 35.4 |
| sgNS ITS+ % | 6.0 |

**Fig S9:** Analysis of fusions/ITS- (ITS-).

A

| | ITS- % | P value |
|---|---|---|
| sgTelo | 4.0 | 0.0790 |
| sgNS | 0.36 | |

B

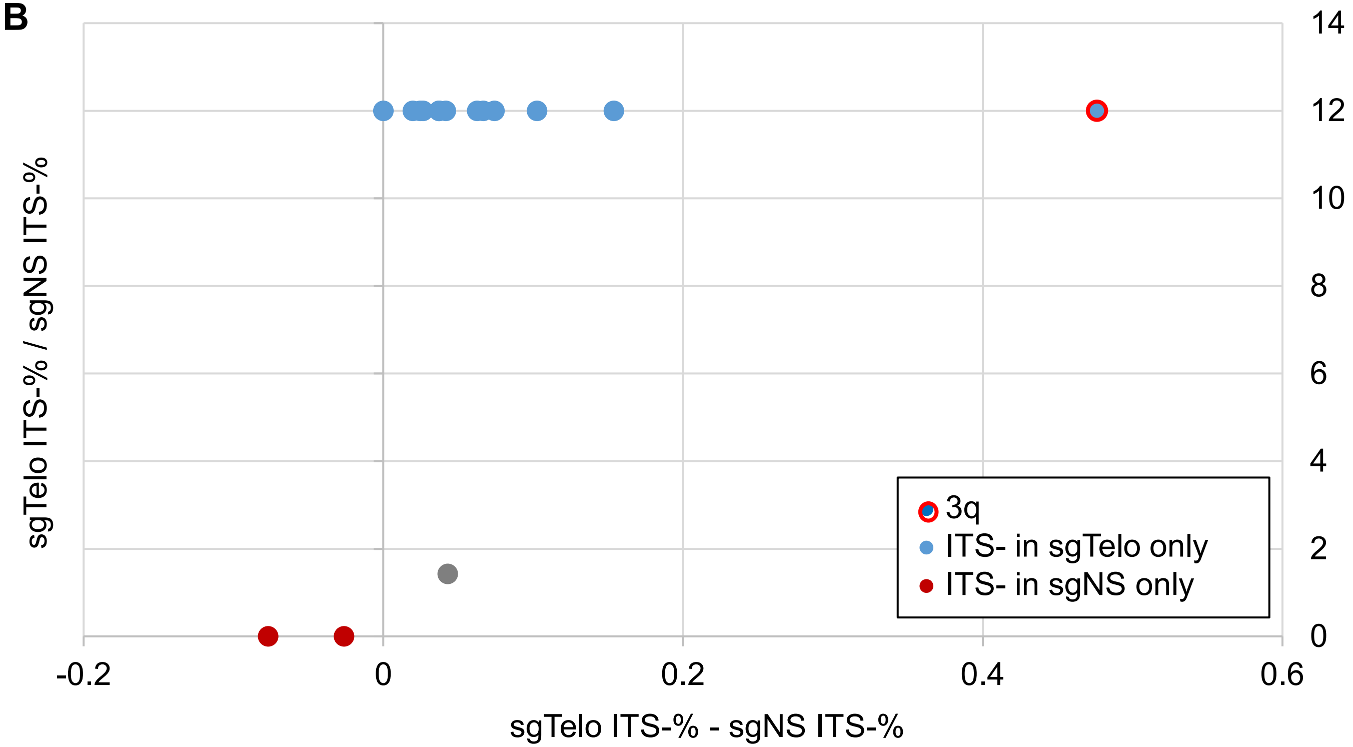


C

| Chromosome arm | 3q |
|---|---|
| sgTelo ITS- % | 47.6 |
| sgNS ITS- % | 0.0 |

**Fig S10:** Analysis of the mean length of telomeres in fusion/ITS+.

**A**

| | Mean length | P value |
|---|---|---|
| sgTelo (kb) | 11.8 | 0.8690 |
| sgNS (kb) | 11.2 | |

**B**

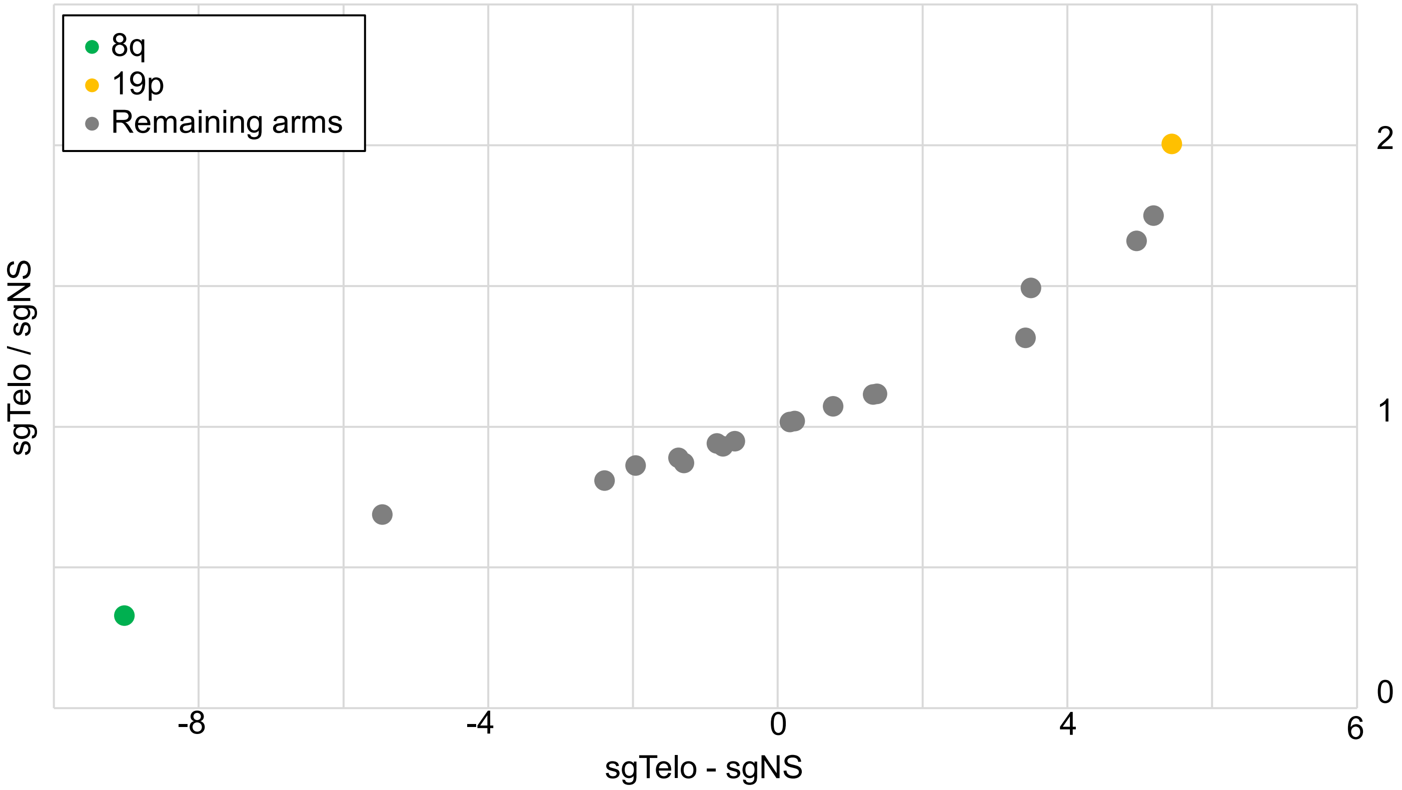


**C**

| Chromosome arm | 8q | 19p |
|---|---|---|
| sgTelo (kb) | 4.4 | 10.9 |
| sgNS (kb) | 13.4 | 5.4 |

**Fig S11**. DNA knots and the staining patterns of 53BP1 on the ITCB

**A**

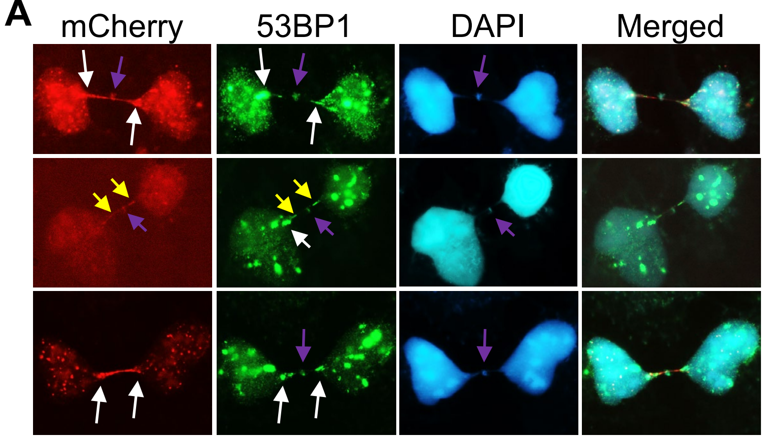


**B**

| | Bridge base | Bridge body | DNA knot |
|---|---|---|---|
| mCherry | 84% | 88% | 72% |
| 53BP1 | 72% | 30% | 70% |
| DAPI | 100% | 56% | 86% |

n = 50 ITCBs

**Fig S12**. EdU pulse labeling assay

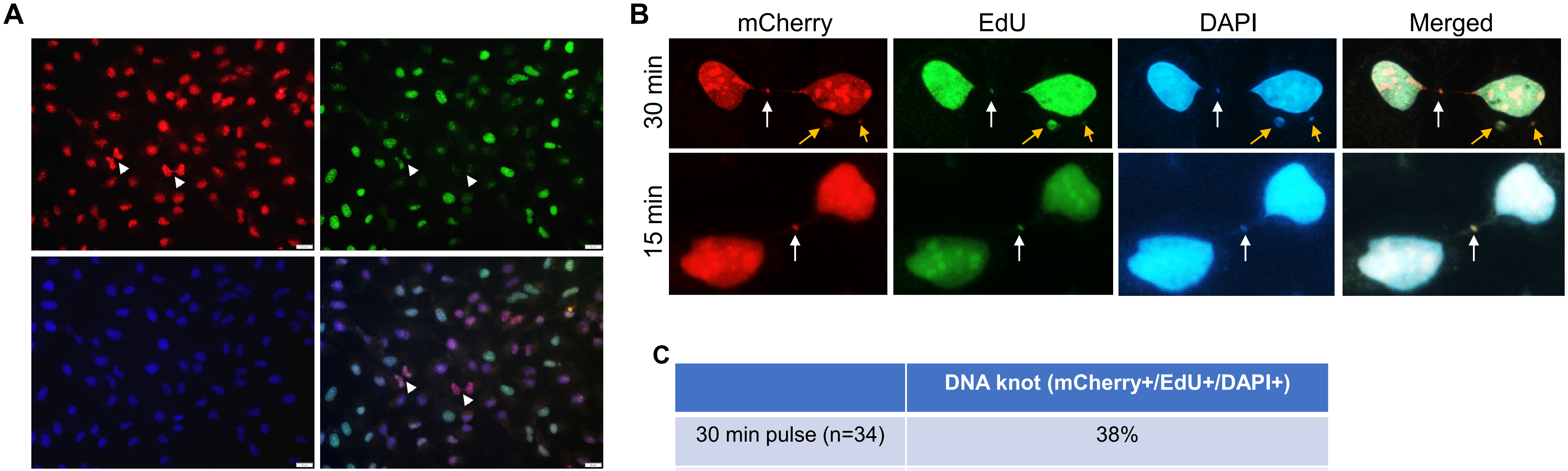


C

| | DNA knot (mCherry+/EdU+/DAPI+) |
|---|---|
| 30 min pulse (n=34) | 38% |
| 15 min pulse (n=32) | 31% |

**Fig S13.** More than half of the DNA knots are positive of Top2A.

**A**

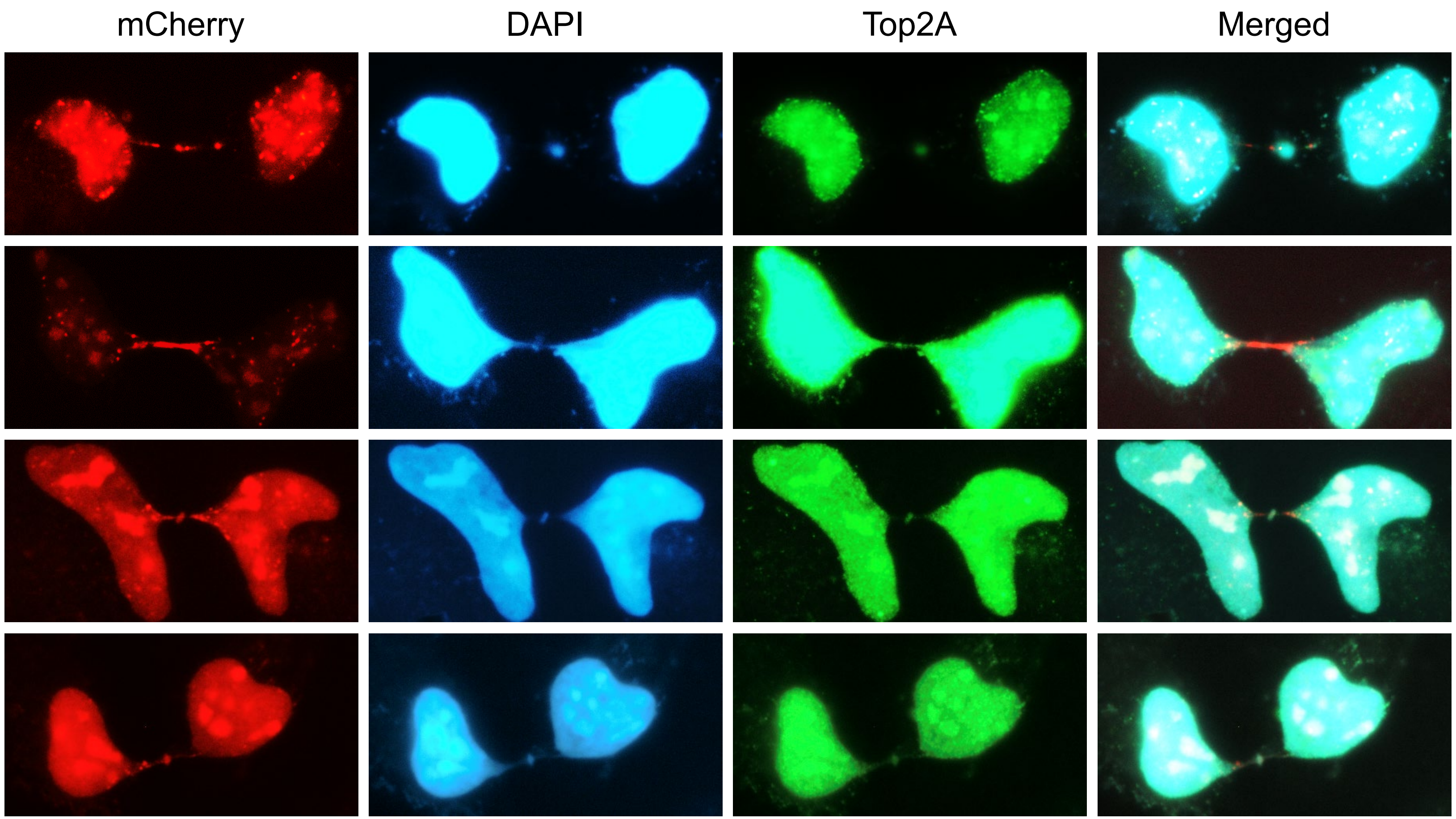


**B**

| % of DNA knots with Top2A | Number of ITCBs |
|---|---|
| 56.6% | >50 |

**Fig S14:** Additional DNA damage response proteins are found at the intercellular telomeric chromosome bridge (ITCB)

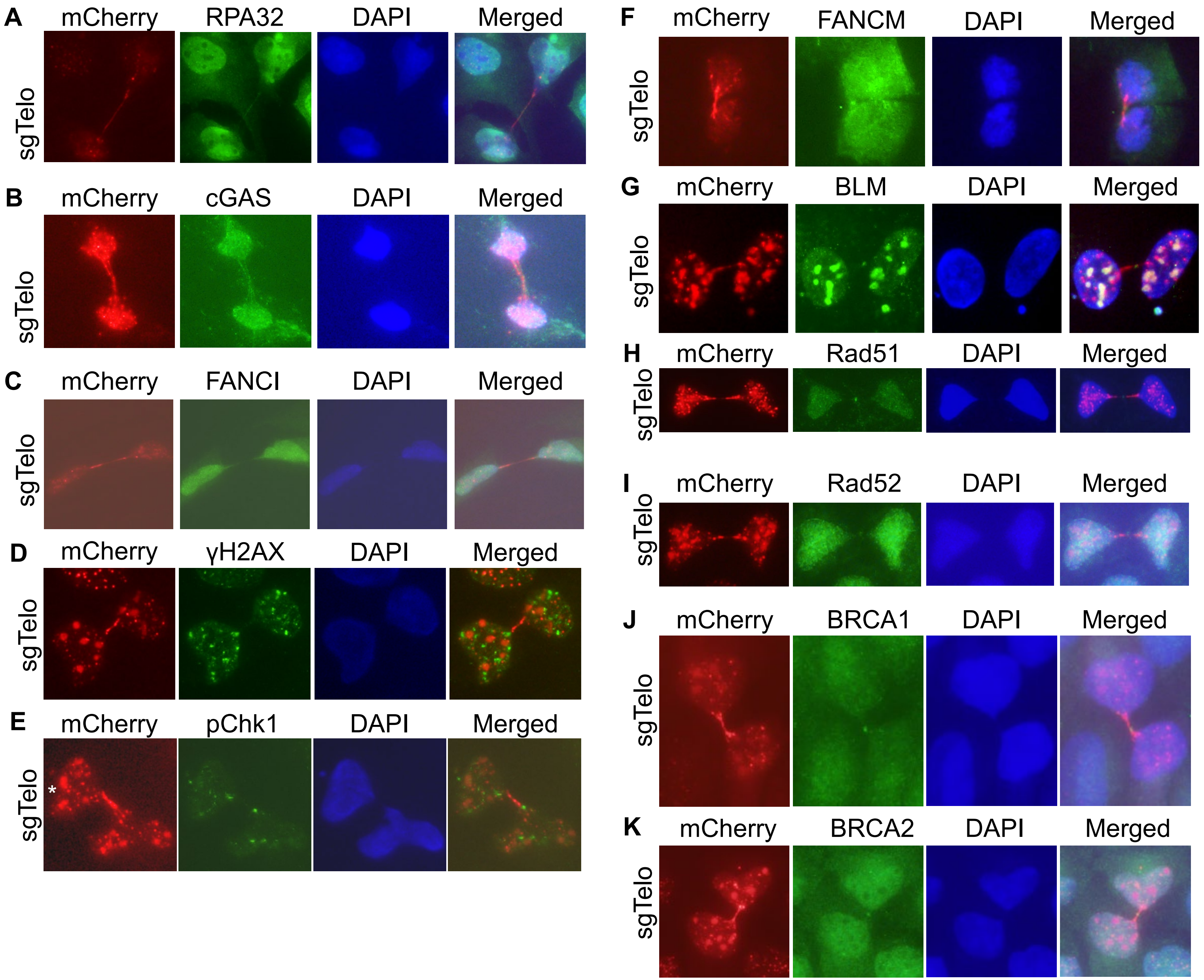


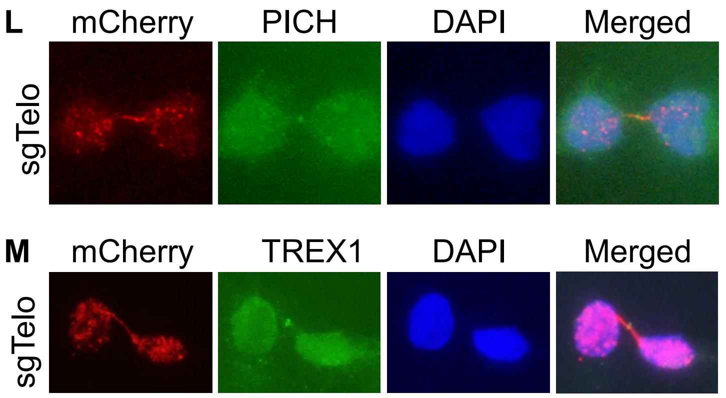


**N**

| **Bridge base** | **53BP1, pChk1, γH2AX** |
|---|---|
| Bridge body | 53BP1, RPA32, cGAS, FANCI, |
| DNA Knot | 53BP1, RPA32, FANCM, BLM, Rad51, Rad52, BRCA1, BRCA2, PICH, TREX1 |

**Fig S15:** Structural features of ITCBs obtained via STORM

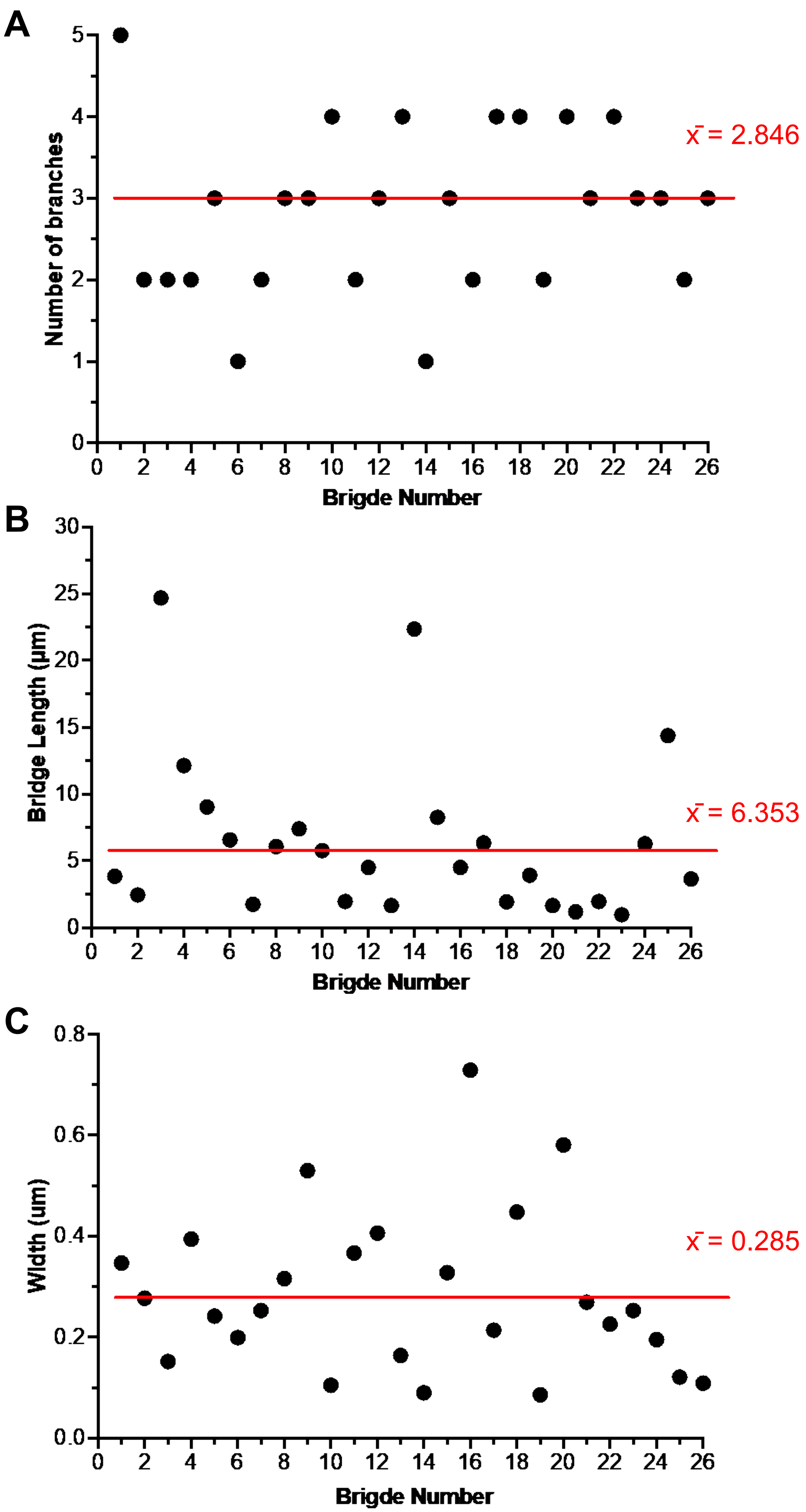

**Fig S16:** Time-lapse analysis of live dCas9/sgTelo cells

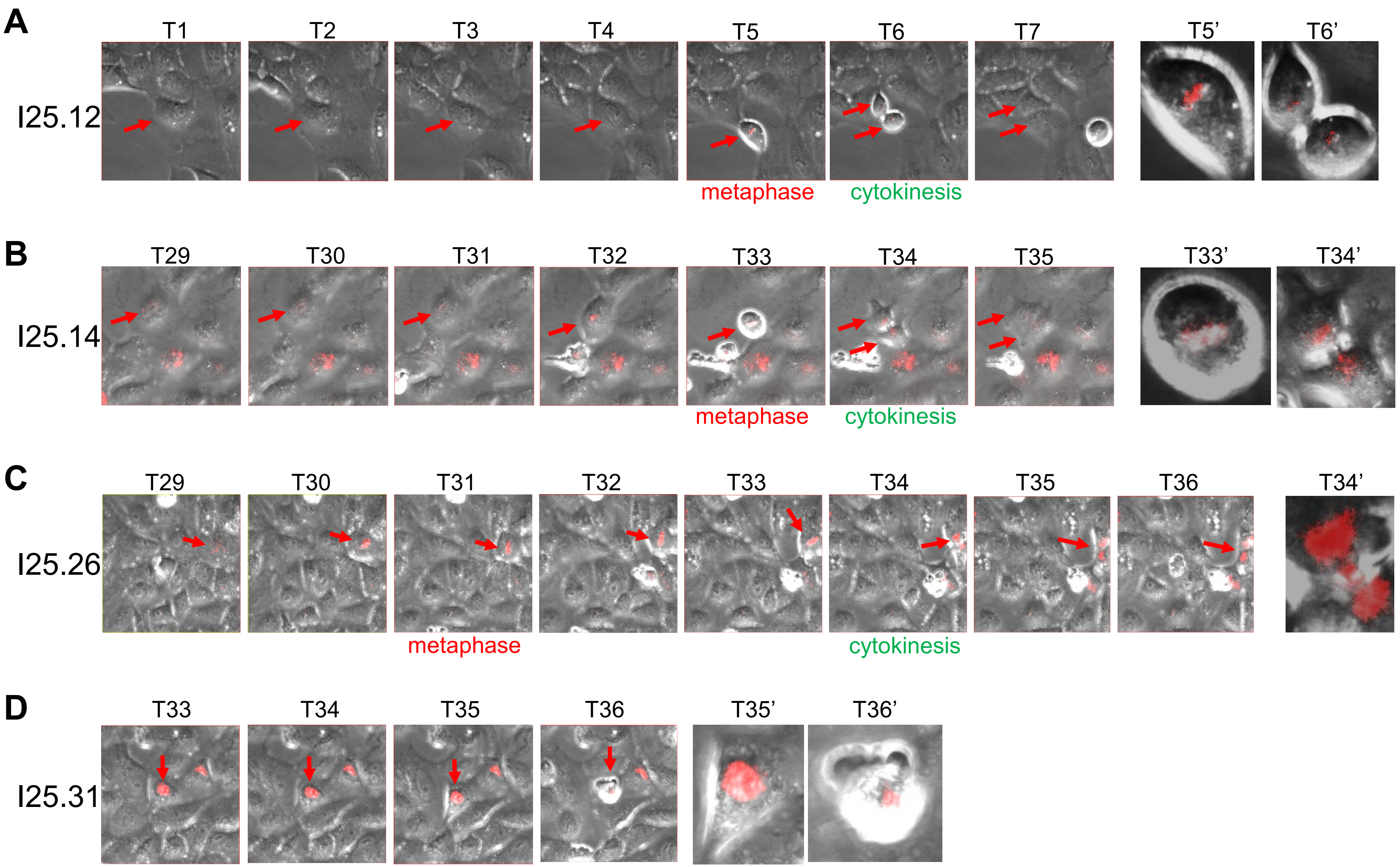

**Fig S17**: depletion of various proteins using siRNA

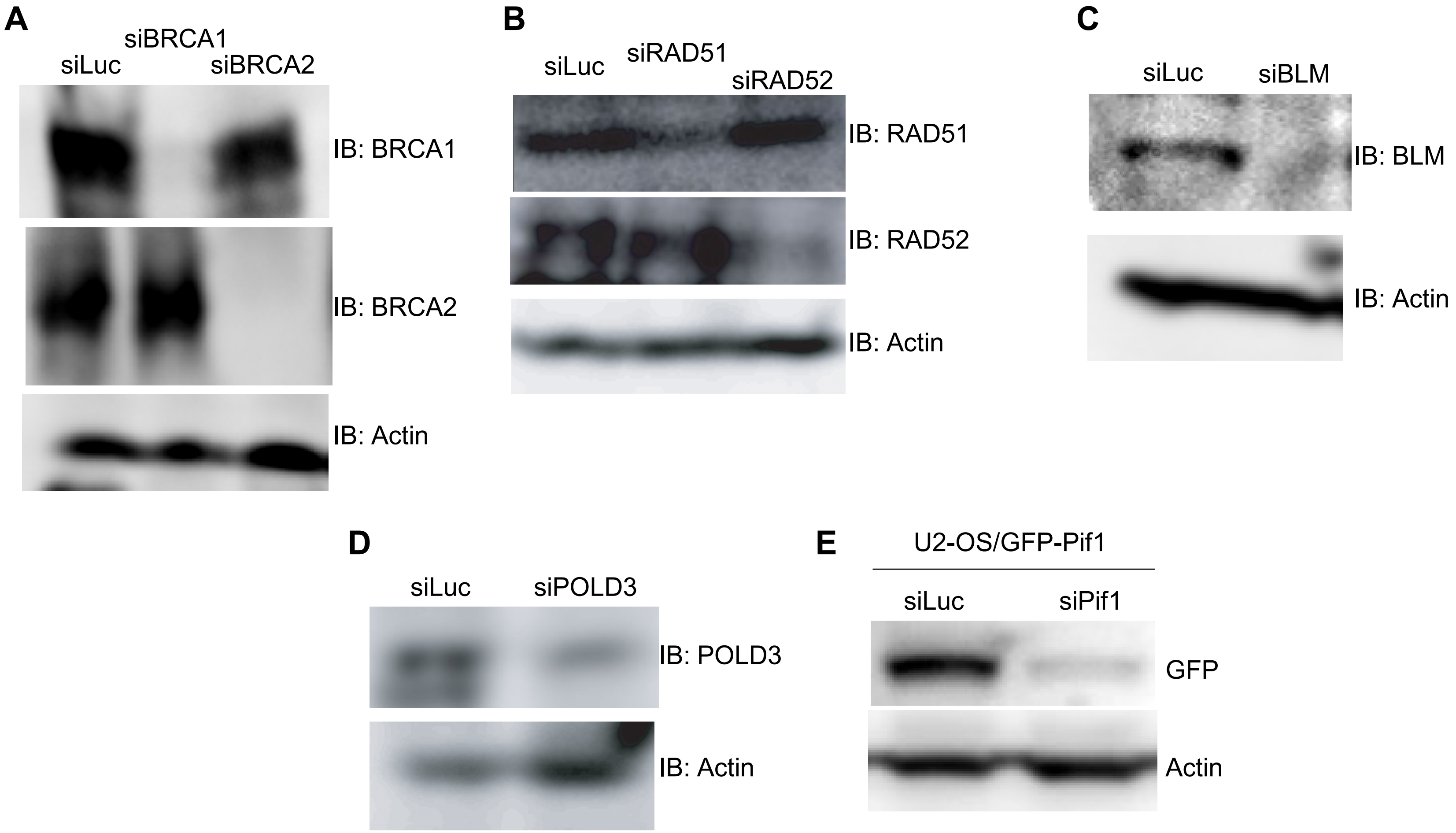

**Fig S18:** Cell cycle analysis of dCas9/sgTelo cells treated with various siRNA and small molecule inhibitors

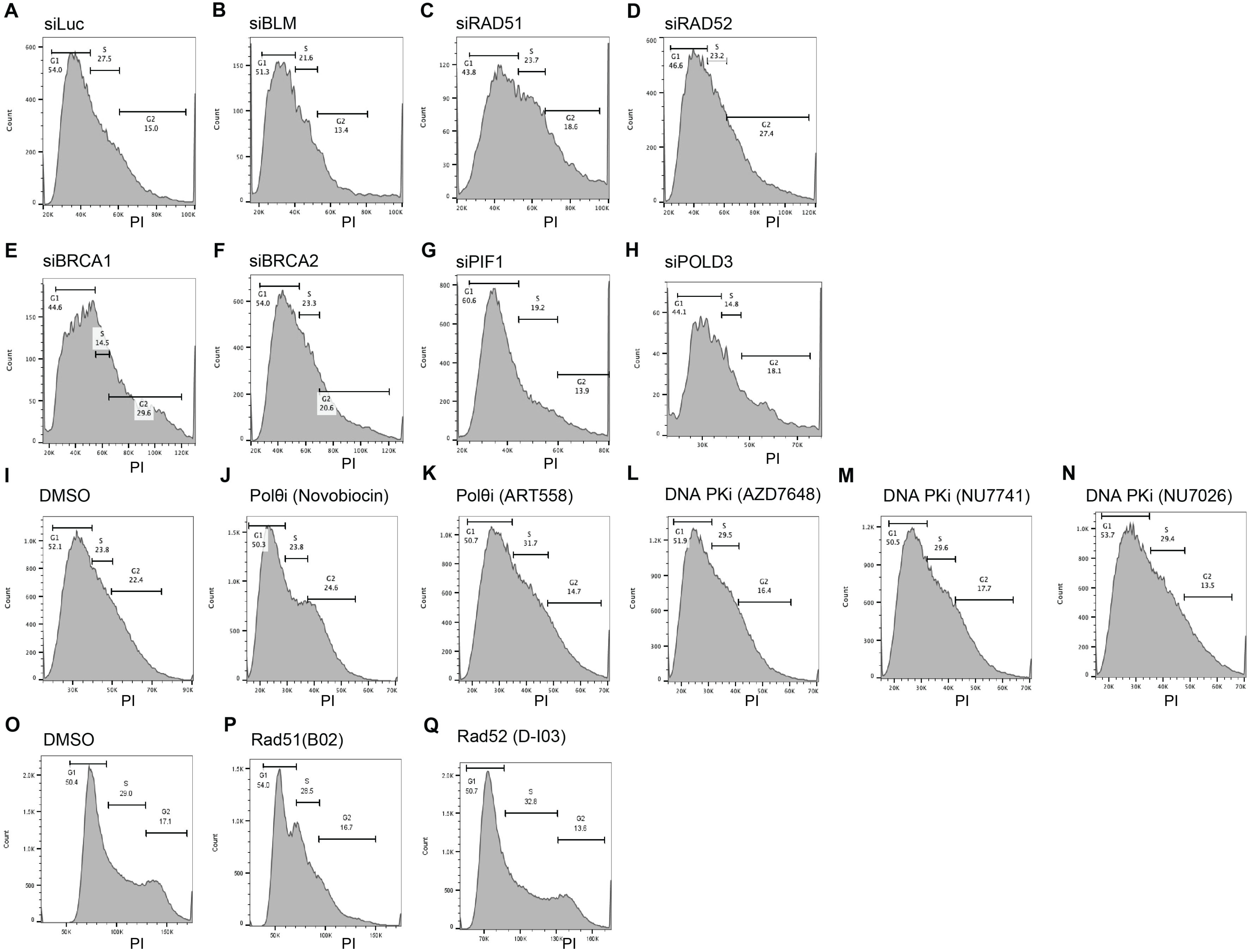
